# Towards the identification of causal genes for age-related macular degeneration

**DOI:** 10.1101/778613

**Authors:** Fei-Fei Cheng, You-Yuan Zhuang, Xin-Ran Wen, Angli Xue, Jian Yang, Zi-Bing Jin

## Abstract

Age-related macular degeneration (AMD) is a leading cause of visual impairment in ageing populations and has no radical treatment or prevention. Although genome-wide association studies (GWAS) have identified many susceptibility loci for AMD, the underlying causal genes remain elusive. Here, we prioritized nine putative causal genes by integrating expression quantitative trait locus (eQTL) data from blood (*n* = 2,765) with AMD GWAS data (16,144 cases vs. 17,832 controls) and replicated six of them using retina eQTL data (*n* = 523). Of the six genes, altering expression of *cnn2*, *sarm1* and *bloc1s1* led to ocular phenotype, impaired vision and retinal pigment epithelium (RPE) loss in zebrafish. Essential photoreceptor and RPE genes were downregulated in *cnn2*- and *sarm1*-knockdown zebrafishes. Through integration of GWAS and eQTL data followed by functional validation, our study reveals potential roles of *CNN2*, *SARM1* and *BLOC1S1* in AMD pathogenesis and demonstrates an efficient platform to prioritise causal genes for human complex diseases.

## Introduction

Age-related macular degeneration (AMD) is an incurable blinding disorder caused by dysfunction of the retinal pigment epithelium (RPE) and progressive loss of photoreceptors in the macula^1^. It results in visual impairment of central vision and disability of daily life activities, such as reading, walking and face recognition. The prevalence of AMD is 8.69% in the age range of 45-85 years globally, and it is projected to affect 196 million people worldwide in 2020^2^. As such, AMD is highly endorsed as a major health and social problem for both individuals and communities, especially in elderly populations^3^.

AMD is one of the most genetically well-defined complex diseases. Genome-wide association studies (GWAS) with increasing sample sizes have identified 52 susceptibility loci which together explain more than 50% of the heritability of liability^4–6^. These findings provide important clues for understanding the genetic architecture of the disease, but the causal genes at those susceptibility loci and underlying mechanisms remain largely unclear. For example, a nonsynonymous variant, *CFH* p. Arg1210Cys (allele frequency = 0.00017 in ExAC), increases AMD risk by >20-fold^7^, but there has been no evidence showing its functional impact on the regulation of gene expression, structural and functional integrity of the protein-coding region, or interplay with the genes nearby^8^. This is partly because of linkage disequilibrium (LD) between single-nucleotide polymorphisms (SNPs) and causative variants that GWAS mapping resolution^9^. This could also be the reason that the trait-associated variants, especially those residing in non-coding regions, exert an impact on gene expression through distal regulation^10^.

With the availability of data from large-scale GWAS^5^, expression quantitative trait locus (QTL) studies^11^ and advanced integrative statistical methods^12,13^, we sought to test the hypothesis that genetic variants at some of the susceptibility loci affect the risk of AMD through genetic regulation of transcriptional levels. We used the summary-data-based Mendelian randomization (SMR) here^12^, which features the statistical power because of the flexibility to utilize GWAS and eQTL data from two independent studies. We subsequently used heterogeneity in dependent instruments (HEIDI) approaches to distinguish causality/pleiotropy (i.e., the same causal variant(s) affecting AMD susceptibility and the expression level of a gene) from linkage (i.e., two distinct causal variants in LD, one affecting AMD susceptibility and the other affecting certain gene expression). This analytical framework has been successfully used in various common diseases, such as diabetes, autoimmune diseases, and psychiatric disorders^14,15^, and is for the first time used in ocular diseases in this study.

We then established an experimental scheme that utilized morpholino oligonucleotide (MO)-induced knockdown and/or mRNA overexpression zebrafish as an animal model to assay the putative causal genes identified by the above integrative analysis (**Figure 1**). Valid procedures were designed to provide morphological and functional assessments of the zebrafish ocular phenotypes, thereby demonstrating the functional relevance to AMD pathogenesis for these prioritised genes. Moreover, the research workflow that combines integrative analysis of large-scale data in humans and functional validation in zebrafish is general and can be used as a new paradigm to efficiently and effectively highlight susceptible genes for human complex diseases and then provide insights for additional prospective therapeutic applications.

**Fig. 1.**
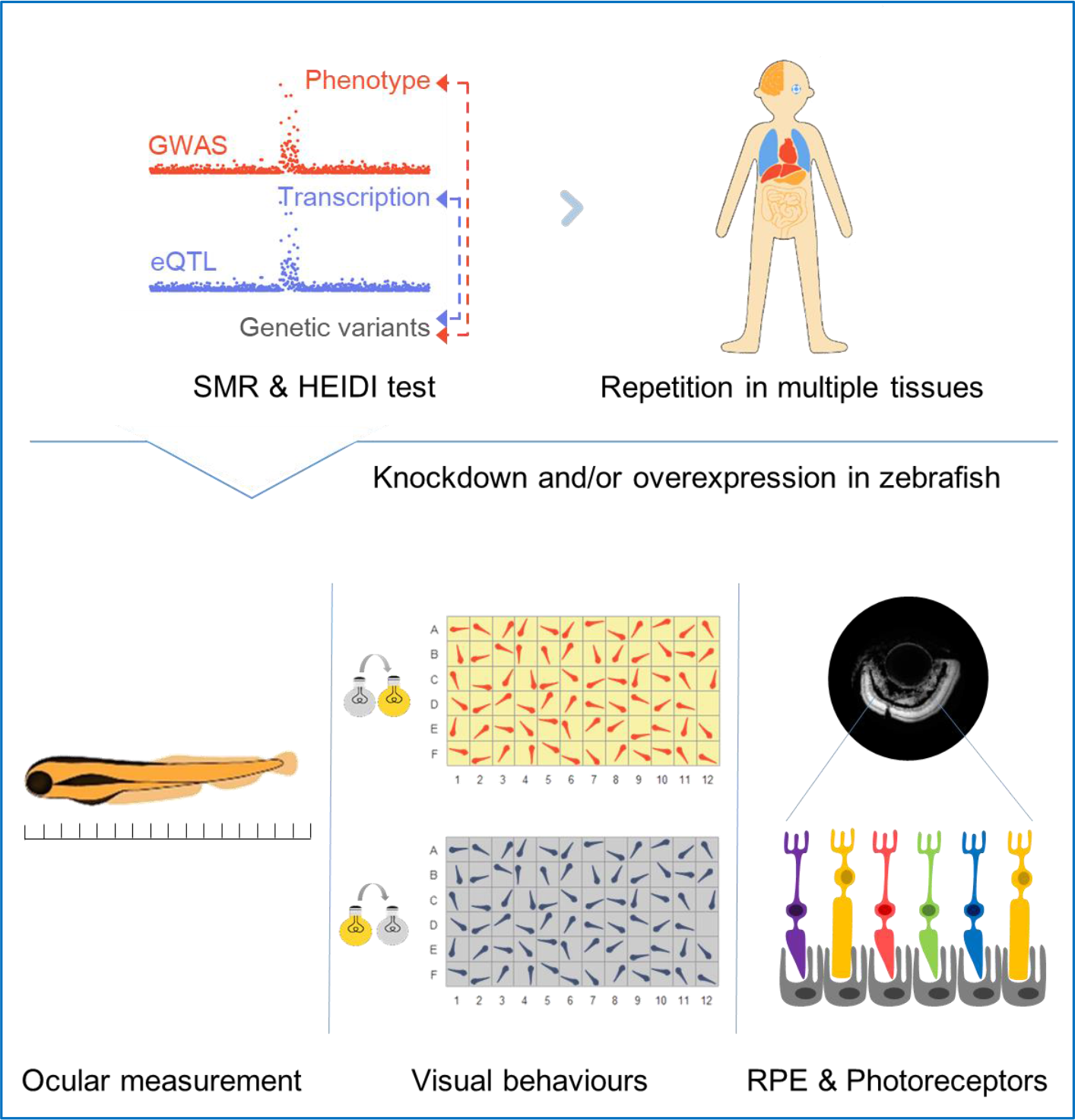
Schematics of study design. First, blood eQTL and AMD GWAS data were integrated through SMR and HEIDI to identify expression–phenotype associations before replications in retina or other 48 human tissues in GTEx. Then, functional experiments were conducted in zebrafish to measure ocular sizes, light-induced behavioural patterns, qPCR of photoreceptors and RPE genes and retinal immunostaining to validate the prioritised genes.

## Results

### Associating gene expression with AMD risk by an integrative analysis

To prioritize genes whose expression levels are associated with AMD risk, we used SMR^12^ to test if a variant has a joint association with AMD risk and the expression level of a gene, using GWAS and eQTL summary data. The GWAS summary data were derived from a study of the International AMD Genomics Consortium with 16,144 AMD cases vs. 17,832 controls^5^. The eQTL summary data were generated by the Consortium for the Architecture of Gene Expression (CAGE) from a study of 36,778 gene expression probes in peripheral blood of 2,765 individuals^11^. All the individuals were predominantly of European ancestry, and both datasets are publicly available (**URLs**). And the test was performed for each of the genes with at least an eQTL at *P*_eQTL_ < 5e−8 (**Methods**).

In total, we identified 16 genes (tagged by 21 probes) at a genome-wide significance level (*P*_*smr*_ = 5.9 ×10^−6^, correcting for 8459 tests, i.e., 8459 probes with at least an eQTL at *P*_eQTL_ < 5e−8) (**Supplementary Table 1**). We then employed the HEIDI method^12^ to reject SMR associations due to linkage (removing probes with *P*_HEIDI_ < 0.05) (**Methods**). And the LD information, required for the HEIDI test, was computed from genotype data of randomly selected 20,000 individuals from European ancestry in the UK Biobank^16^. Consequently, 9 genes (tagged by 12 probes) were retained (**Table 1**), and some of which, including *BLOC1S1*, *PILRB*, and *TMEM199*, have been reported in a recent study that used a different strategy to integrate AMD GWAS with retinal eQTL data^17^. We found that 58.3% of the identified probes were not tagging the closest genes to the top associated GWAS signals, consistent with the observations from previous studies^10,12^. The eQTL variants of all the prioritised genes were common with minor allele frequencies (MAF) ranging from 0.11 to 0.49. It is of note that the association of rs7212349 (i.e., the top associated variant for gene *SARM1* with *P*_eQTL_ = 1.0e−22) with AMD (*P*_GWAS_ = 1.8e−7) did not reach the conventional genome-wide significance threshold, suggesting a gain of power in gene discovery by leveraging eQTL data, in line with previous work^12^.

**Table 1.**
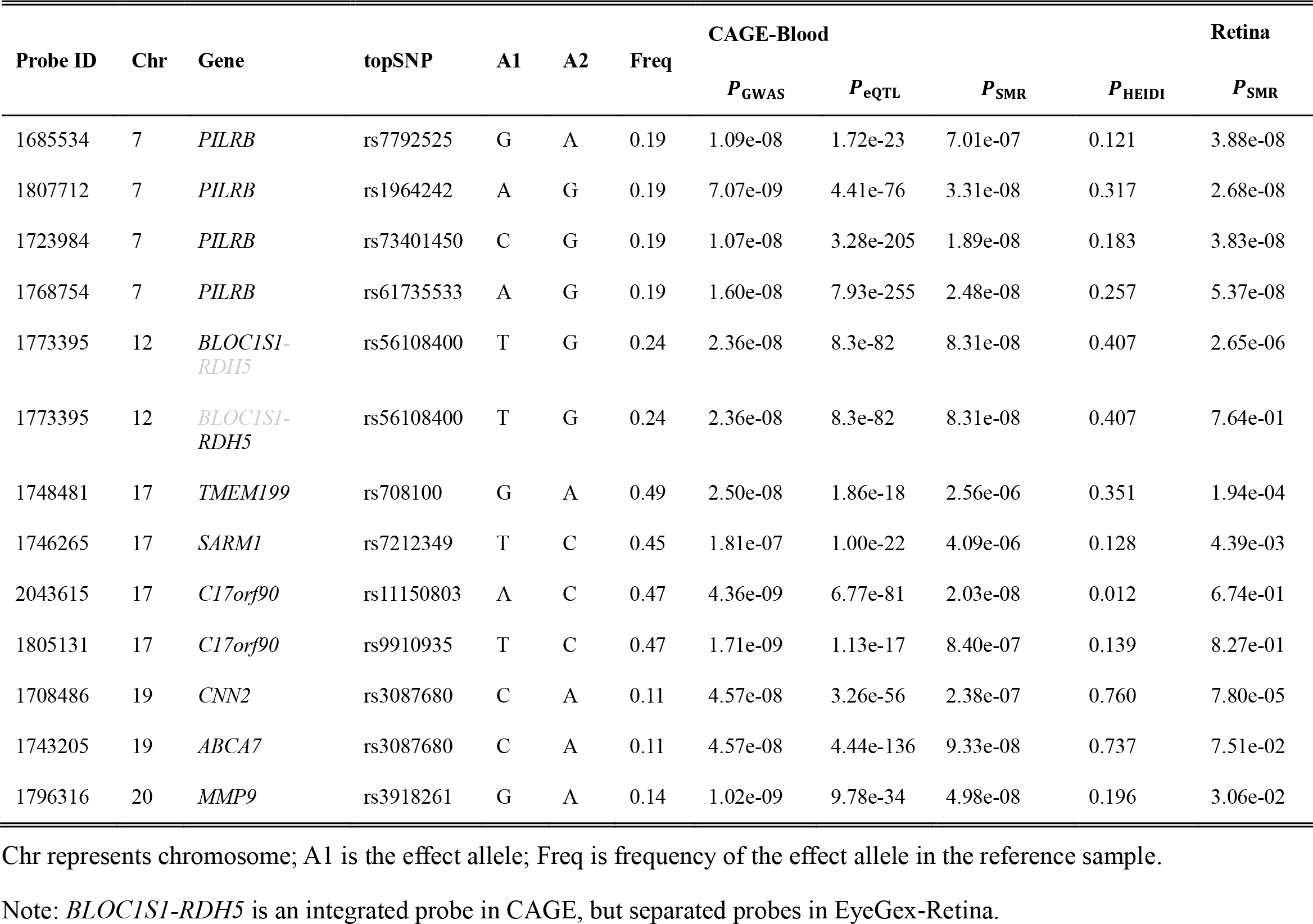
Putative causal genes for AMD identified from SMR & HEIDI analysis using the blood or retina eQTL data.

| Probe ID | Chr | Gene | topSNP | A1 | A2 | Freq | CAGE-Blood |  |  |  | Retina |
| --- | --- | --- | --- | --- | --- | --- | --- | --- | --- | --- | --- |
| | | | | | | | $P_{\text{GWAS}}$ | $P_{\text{eQTL}}$ | $P_{\text{SMR}}$ | $P_{\text{HEIDI}}$ | $P_{\text{SMR}}$ |
| 1685534 | 7 | PILRB | rs7792525 | G | A | 0.19 | 1.09e-08 | 1.72e-23 | 7.01e-07 | 0.121 | 3.88e-08 |
| 1807712 | 7 | PILRB | rs1964242 | A | G | 0.19 | 7.07e-09 | 4.41e-76 | 3.31e-08 | 0.317 | 2.68e-08 |
| 1723984 | 7 | PILRB | rs73401450 | C | G | 0.19 | 1.07e-08 | 3.28e-205 | 1.89e-08 | 0.183 | 3.83e-08 |
| 1768754 | 7 | PILRB | rs61735533 | A | G | 0.19 | 1.60e-08 | 7.93e-255 | 2.48e-08 | 0.257 | 5.37e-08 |
| 1773395 | 12 | BLOC1S1-RDH5 | rs56108400 | T | G | 0.24 | 2.36e-08 | 8.3e-82 | 8.31e-08 | 0.407 | 2.65e-06 |
| 1773395 | 12 | BLOC1S1-RDH5 | rs56108400 | T | G | 0.24 | 2.36e-08 | 8.3e-82 | 8.31e-08 | 0.407 | 7.64e-01 |
| 1748481 | 17 | TMEM199 | rs708100 | G | A | 0.49 | 2.50e-08 | 1.86e-18 | 2.56e-06 | 0.351 | 1.94e-04 |
| 1746265 | 17 | SARM1 | rs7212349 | T | C | 0.45 | 1.81e-07 | 1.00e-22 | 4.09e-06 | 0.128 | 4.39e-03 |
| 2043615 | 17 | C17orf90 | rs11150803 | A | C | 0.47 | 4.36e-09 | 6.77e-81 | 2.03e-08 | 0.012 | 6.74e-01 |
| 1805131 | 17 | C17orf90 | rs9910935 | T | C | 0.47 | 1.71e-09 | 1.13e-17 | 8.40e-07 | 0.139 | 8.27e-01 |
| 1708486 | 19 | CNN2 | rs3087680 | C | A | 0.11 | 4.57e-08 | 3.26e-56 | 2.38e-07 | 0.760 | 7.80e-05 |
| 1743205 | 19 | ABCA7 | rs3087680 | C | A | 0.11 | 4.57e-08 | 4.44e-136 | 9.33e-08 | 0.737 | 7.51e-02 |
| 1796316 | 20 | MMP9 | rs3918261 | G | A | 0.14 | 1.02e-09 | 9.78e-34 | 4.98e-08 | 0.196 | 3.06e-02 |

Chr represents chromosome; A1 is the effect allele; Freq is frequency of the effect allele in the reference sample.
Note: *BLOC1S1-RDH5* is an integrated probe in CAGE, but separated probes in EyeGex-Retina.

### Replication of the SMR associations in retina and other tissues

Given possible tissue-specific genetic effects, we replicated SMR associations of the nine significant genes in retina. The retinal eQTL summary data were retained from 523 postmortem subjects^17^, available in the Eye Genotype Expression (EyeGex) database (**URLs**). Six of the nine genes were replicated at *P*_*smr*_ = 5.6 × 10^−3^(i.e., 0.05/9), a relatively high replication rate given the small sample size of the EyeGex data (**Table 1** and **Supplementary Figure 1**). However, for some of the replicated genes, the eQTL effects were significantly different, even in opposite directions, between the retina and CAGE-blood. For instance, the estimated effect of *SARM1* on AMD risk was 0.301 (standard error (SE) = 0.065 and *P*_eQTL_ = 4.1e−06) in CAGE-blood, consistent in several other tissues (see **Supplementary Figure 1**), but was −0.351 (SE = 0.123 and *P*_eQTL_ = 4.4e−03) in retina, suggesting that a strong tissue-specific effect and the importance to replicate and interpret discovery results in disease-relevant tissue(s).

It is unclear whether AMD is a localized disease (occurring only in affected retinas) or an ocular manifestation of a systemic process, since risk factors such as cigarette smoking, nutrition, and cardiovascular disease have a significant impact on disease progression^19–23^. Thus, we conducted SMR analysis for the nine genes in a wider range of tissues available in the GTEx project (**Supplementary Figure 1**). Intriguingly, the results showed that *PILRB*, a key activator in immune function, is significant across all 48 human tissues with effect sizes ranging from 0.081 (in retina) to 0.249 (in whole blood), implying that systemic immune pathways may be involved in AMD pathogenesis. In fact, *PILRB* is not an exception; all the nine genes were significant in at least two tissues and did not display obvious tissue-specific effects except for *SARM1* shown above, consistent with the results from previous studies that cis-eQTL effects are largely consistent across tissues^24^. Our results also suggest that the across-tissue replication rate depends heavily on the size of replication sample, supporting that using blood eQTL data from large samples gains power for gene discovery.

### Knockdown of *cnn2* or *sarm1* led to ocular abnormalities in zebrafish

To validate the functional relevance of the prioritised genes to AMD pathogenesis, we sought an animal model that is amenable to manipulating gene expression and that can reliably evaluate ocular phenotypes. We chose zebrafish also because it has been extensively used to model ocular and other disorders^25–27^. Accounting for sequence homology between human and zebrafish, we obtained 4 of the 9 prioritised genes (*CNN2*, *SARM1*, *BLOC1S1* replicated at *P*_SMR_ < 5.6e−3 and *MMP9* at *P*_SMR_ < 0.05 using the retina eQTL data; **Table 1**) with orthologue similarity above 60%, reported by either Ensembl or GeneCards, for functional follow-up (**URLs, Supplementary Table 2**). Then, MO technology, blocking the translation process of a certain mRNA, was used to knock down the corresponding gene. We observed that downregulation of *cnn2* and *sarm1* at doses of 6 ng MO and 1.0 ng, respectively, led to obvious decreases in axis length (by 32.5% and 21.3%, respectively) and eye area (by 48.5% and 28.5%, respectively), whereas suppression of *mmp9* and *bloc1s1* showed no significant difference at 3 days post fertilisation (dpf) (**Figure 2** and **Supplementary Figure 2**).

**Fig. 2.**
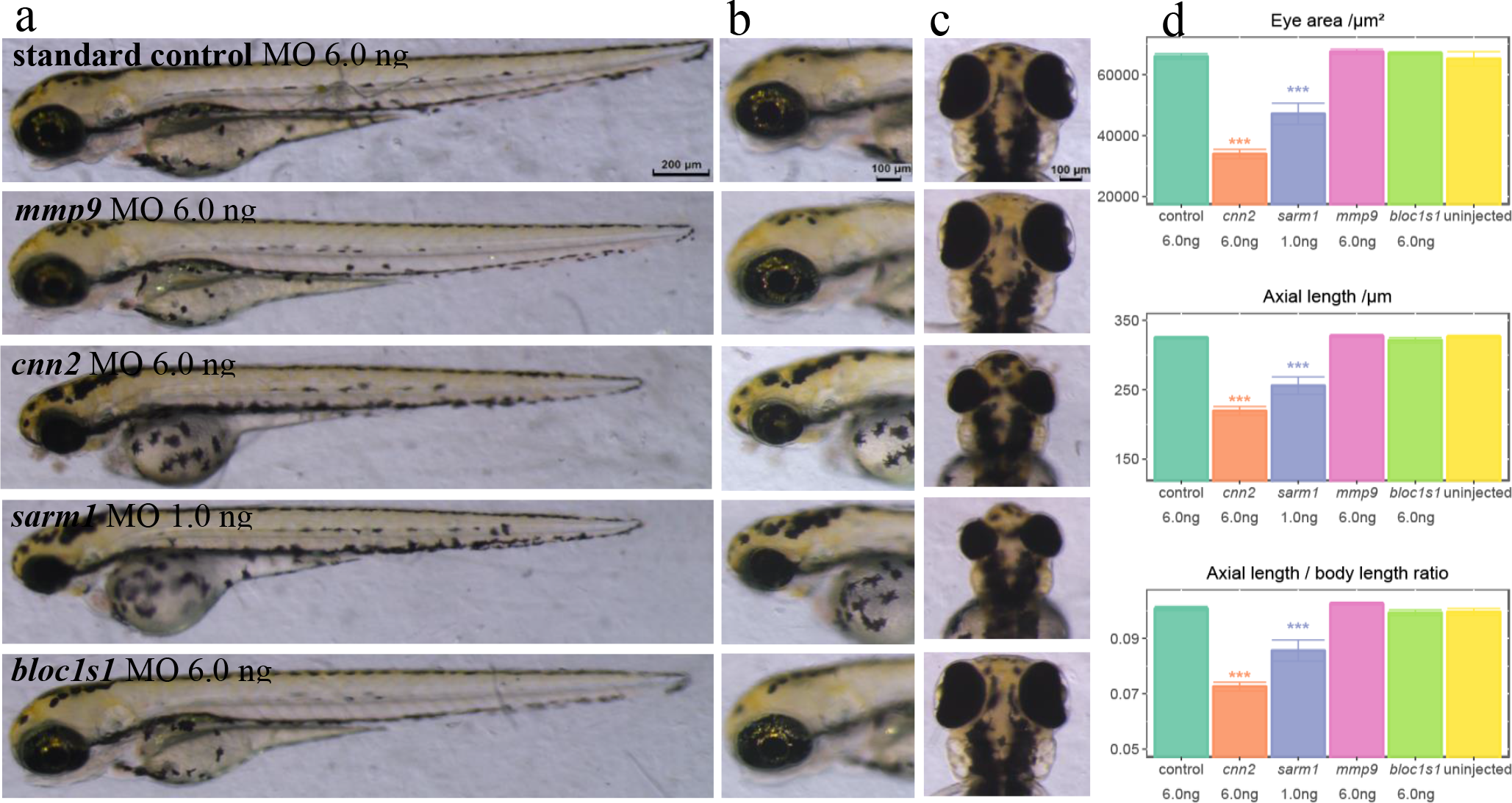
Phenotypes of cnn2-, sarm1-, mmp9-, bloc1s1-deficient zebrafish. (**a**) Lateral view of whole bodies. (**b**) Magnified lateral view of zebrafish eyeballs exhibits apparent decreases in eye area for *cnn2*- and *sarm1*-deficient fishes. (c) Vertical view of eyeballs showing shorter axial length in *cnn2* and sarm1 knockdown larvae. (d) Quantification of eye area, axial length and ratio of axial length and body length, respectively. Bar plots are shown as the mean±s.e.m. T-test was performed between each group and the standard control. *P<0.05, **P<0.01, ***P<0.001. N=10 for each group.

To further confirm the results, we conducted dose-dependent and rescue experiments. We found that in comparison to the control, eye sizes of *cnn2*-MO morphants were smaller and the degree was inversely proportional to the MO dose (from 4.0 to 6.0 ng) without higher mortality (**Supplementary Figure 3**). For the *sarm1*-MO morphants, abnormal ocular phenotypes occurred when the MO dose was extremely low (0.50 ng), and the impact accumulated when the dose increased (from 0.50 to 4.0 ng), indicating that ocular development might be sensitive to the *sarm1* expression level (**Supplementary Figure 4**). Importantly, rescue of the smaller eyes was achieved in the mRNA and MO co-injected larvae for both genes (**Figure 3a**,**d**). Axial length was recovered by 95.8% for *sarm1*-MO and 91.0% for *cnn2*-MO (**Supplementary Note 1**). This study provides evidence for potential roles of *CNN2* and *SARM1* in ocular development and disease-causing mechanisms.

**Fig. 3.**
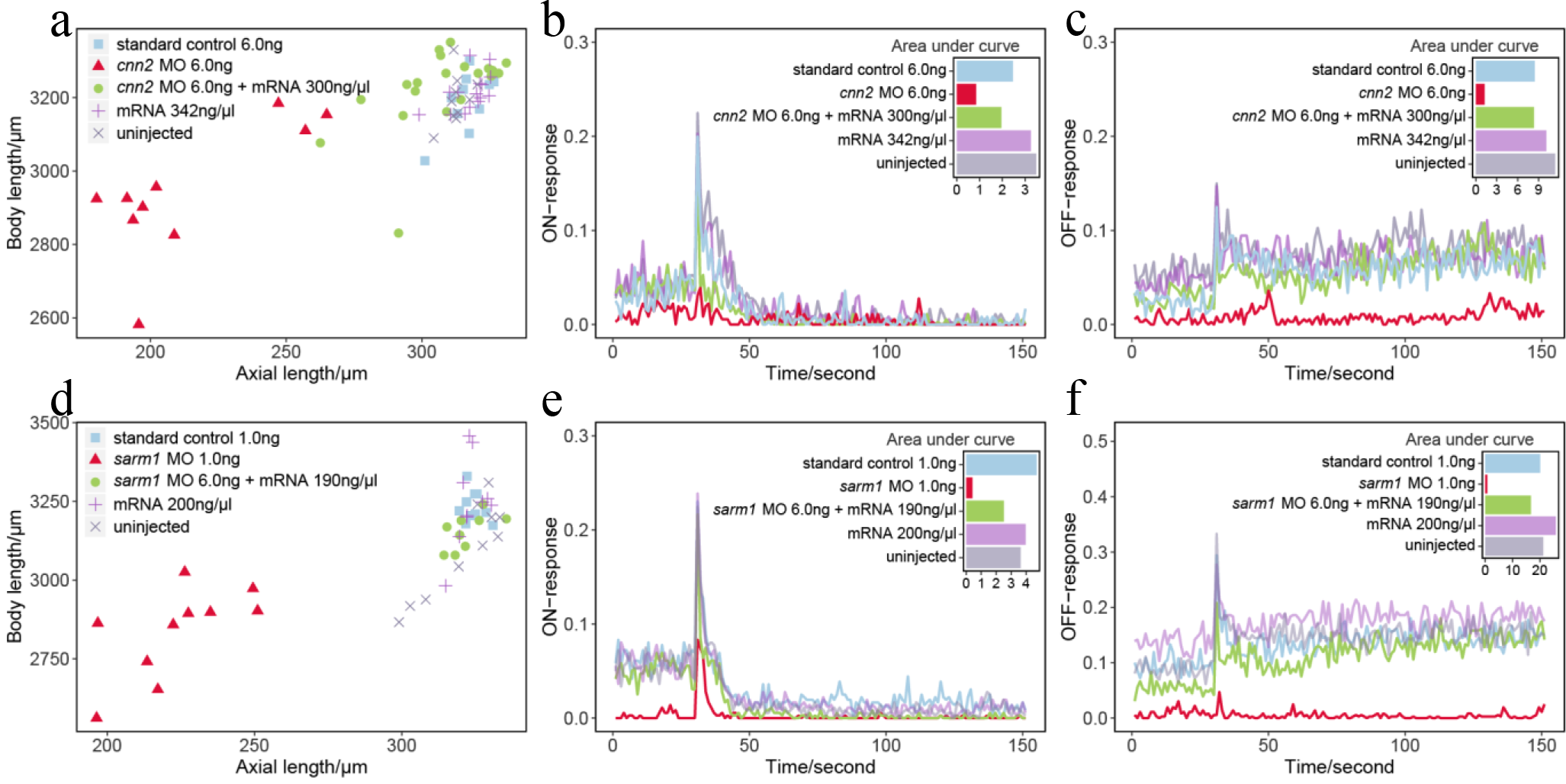
Morphological and functional effects of knockdown and rescue of *cnn2* and *sarm1* in zebrafish larvae. (**a, d**) The scatter plots of body length and eyeball axial length. The *cnn2* or *sarm1* MO group (labelled by red triangle) was clearly separate from the other groups, showing a reduction in eye size. In addition, injecting corresponding mRNA (labelled by green dot) could rescue the reduced ocular size. (**b, c, e, f**) VMR testing for ON and OFF responses. Lights-ON or lights-OFF stimuli occurred at 30 s. Real-time motor activities of zebrafish were recorded by lines. The area under the curve reflected the sum of motor activities during the 150 s. The responses of the *cnn2* or *sarm1* MO group (labelled by red line) were dramatically weakened and could also be saved by injecting mRNA (labelled by green line).

### Knockdown of *cnn2* or *sarm1* led to functional impairment in zebrafish

We then used a visual motor response (VMR) assay to evaluate the visual condition at the behavioural level 5 dpf. According to a standard protocol^28^, zebrafish were placed in a 96-well plate and the locomotor response to light alteration was monitored. We found that, for *CNN2*, three groups, including the uninjected control, standard MO control and *cnn2* mRNA, had a brief spike at approximately 0.20-0.22 of motor activity for ON response and 0.12-0.15 for OFF response (**Figure 3b, 3c**). In comparison, the response of the *cnn2* MO-injected group was weakened and delayed, with the peak dramatically decreasing by 61.9% for lights-ON and by 78.6% at 20 s later when lights-OFF. Of note, augmentation of *cnn2* mRNA with MO morphants could save a visual response to 0.16 for lights-ON (recovering by 61.5%); when lights-OFF, their motor activities were intensified to 0.08 (recovering by 45.5%), and the baseline was notably improved. Actually, functional recovery relied on a sufficient dose of injected mRNA. Our pre-experiment suggested that only partial rescue of visual function could be realized when injecting less *cnn2* mRNA (**Supplementary Figure 5**). In addition, the *SARM1* group showed similar results but with more severe visual impairment (**Figure 3e,f**). Notably, simple injection of *sarm1* mRNA seemed to slightly promote the OFF response, implying an underlying therapeutic target. Taken together, the results of the visual function assays further elucidate the reduction in visual motor activities specifically caused by forced downregulation of *cnn2* or *sarm1*, indicating their essential roles in maintaining normal visual function.

### Knockdown of *cnn2* or *sarm1* downregulated the expression of photoreceptor and RPE signature genes

Considering the potential role of *CNN2* and *SARM1* in AMD pathophysiology, we sought to test the hypothesis whether visual impairment could be attributed to defects of vital photoreceptor or RPE genes. Thus, we conducted reverse transcription quantitative polymerase chain reaction (RT-qPCR) analysis to measure the expression levels of retinal signature genes: *rhodopsin*, four kinds of cone opsins (*red*, *green*, *blue*, and *uv*) and RPE-specific gene *rpe65a*^29,30^. For the *cnn2* 6.0 ng MO group and the *sarm1* 1.0 ng MO group, all these genes showed dramatic decreases compared to the standard MO control; *rhodopsin* decreased by 117.1-fold and 12.6-fold respectively, so as to cone opsin (*green opsin* by 797.5-fold and 13.6-fold, *blue opsin* by 279.8-fold and 10.4-fold, *red opsin* by 107.4-fold and 8.0-fold, and *uv opsin* by 19.2-fold and 10.8-fold), and *rpe65a* (by 7.5-fold and 3.6-fold) (**Figure 4**). However, regarding the *mmp9* 6.0 ng MO group, the expression levels of all six genes slightly increased by 1.2-2.4 folds. Combined with the results above, the transcriptional reduction of signature genes in RPE and photoreceptors, caused by downregulation of *cnn2* or *sarm1*, is in line with ocular abnormality and visual behaviour impairments. Since opsin gene expression is a determinant factor for photoreceptor degeneration^31–33^, we hypothesize that downregulation of *CNN2* and *SARM1* are likely involved in photoreceptor degeneration during the pathogenic process of AMD.

**Fig. 4.**
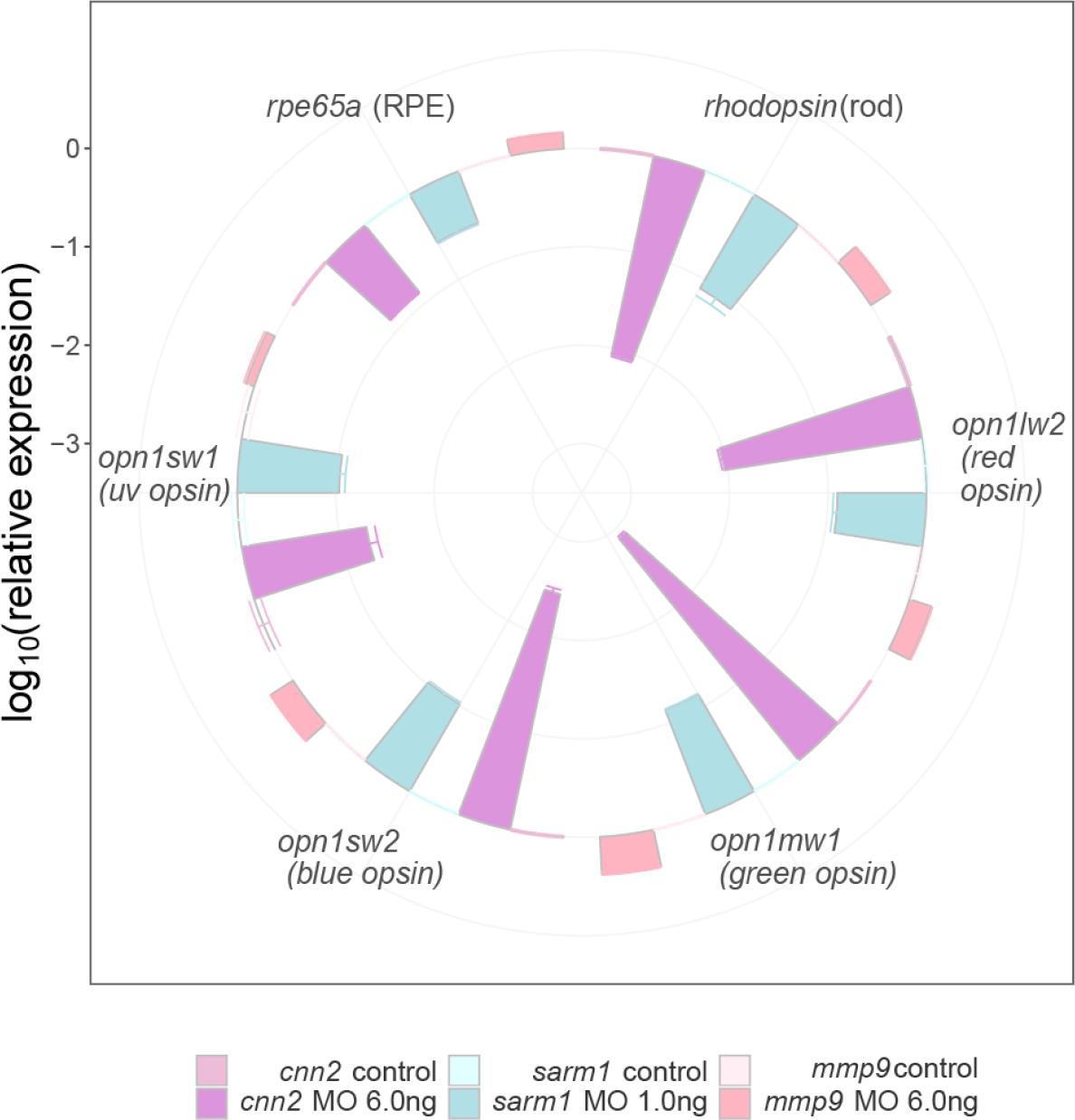
Real-time qPCR of photoreceptor and RPE genes in the MO zebrafish oculus. The x axis represents photoreceptor and RPE signature genes, and the y axis represents the *log*_10_ (*relative expression*). Centripetal bars indicate downregulation, and centrifugal bars indicate upregulation. The *cnn2*- and *sarm1*- MO groups revealed relatively low expression for all gene markers, whereas the *mmp9* group showed a slight increase. The bar plot is shown as the mean±s.e.m.

### Knockdown of *cnn2* or *sarm1* disrupted RPE in zebrafish

In addition to the alteration of photoreceptors and RPE at transcriptional level, we investigated the retinal morphological consequences of the knockdown by cryosection and immunostaining to further verify the functional relevance of *CNN2* and *SARM1* with AMD pathogenesis. Using DAPI for nuclear staining, zpr-1 for cones, zpr-2 for RPE and zpr-3 for rods, we found that laminations were basically intact in both the *cnn2* 6.0 ng MO group and the *sarm1* 1.0 ng MO group (**Figure 5**). For the *cnn2* 6.0 ng MO group, staining of cones and rods showed no apparent difference from the standard control, but RPE displayed obvious disorganization and cell loss. For the *sarm1* 1.0 ng MO group, staining of RPE also demonstrated significant deficiency, with most rods disrupted (**Supplementary Figure 6**), whereas cones remained complete. In fact, a prior experiment of the *sarm1* 6.0 ng MO group observed severe lamination disruption, indicating that extreme insufficiency of *sarm1* has a severely adverse impact on ocular development (**Supplementary Figure 7**). Altogether, for both *cnn2* and *sarm1*, RPE were morphologically injured, which is in accordance with a well-accepted understanding that degeneration of RPE is a fundamental trigger of the cascade of events resulting in AMD pathology^34^. However, photoreceptors seemed not significantly affected. A likely hypothesis is that, in a certain condition, functional changes might precede morphological changes. That is, in the developmental stage, a small number of healthy RPE tissues could burden delivering oxygen and metabolites, where photoreceptors maintain morphological normality but are gradually functionally injured due to downregulation of *cnn2* or *sarm1* and photoreceptor core genes, then cumulative damage tips the balance and consequently leads to degeneration^34^.

**Fig. 5.**
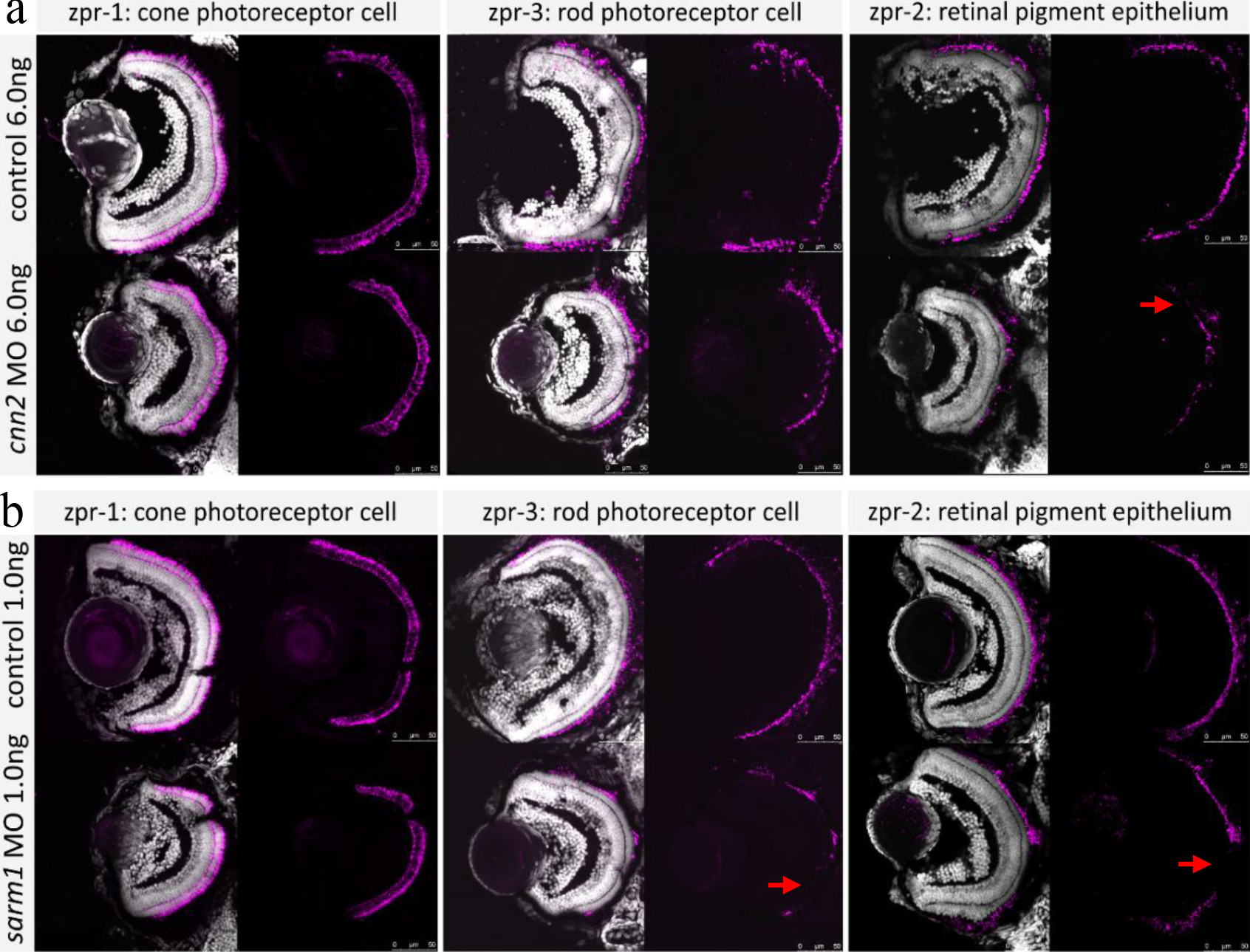
Retinal architecture of *cnn2*- and *sarm1*-deficient morphants. Nuclear layers were stained by DAPI in grey, and each group showed intact lamination. zpr-1, zpr-2 and zpr-3 were stained in purple for each column. (**a**) Signals of zpr-1 and zpr-3 in the *cnn2* MO 6.0 ng group were detected at relatively normal levels, but the signal of zpr-2 was much lower compared with the standard control. (**b**) Signals of zpr-3 and zpr-2 were distributed significantly less in the out layer of retina for the *sarm1* MO 1.0 ng group, and zpr-1 remained intact.

### Overexpression of *bloc1s1* resulted in mild impaired ocular phenotypes in zebrafish

Given the significant SMR associations of *BLOC1S1* in both discovery and replication analyses where the effect size was positive in retina and negative in blood, we assumed that GWAS loci in *BLOC1S1* affect the risk of AMD by upregulating its expression. Therefore, we injected an additional zebrafish group with *bloc1s1* mRNA. Compared to controls, smaller eyes were observed in *bloc1s1*-overexpressing fishes (**Figure 6a**) with axial length decreased by 7%. For the visual behaviour, the ON response was not apparently affected, but impairment of the OFF response was observed in the *bloc1s1* mRNA-injected group with the peak of OFF response declined by 59.1% (**Figure 6b,c**). Immunostaining results showed that disruption of cones, rods and RPE in *bloc1s1*-overexpression group occurred (**Figure 6d**) but more than one half of staining repetition had no notable difference in comparison to the standard control (**Supplementary Figure 8**). In terms of qPCR, however, rod gene was down-regulated by 1.4-fold (*P* = 5.4e−4) and uv cone gene was up-regulated significantly by 1.3-fold (*P* = 0.025) but other retinal genes remained same (**Supplementary Figure 9**), likely in the early stage of degeneration since a previous study showed that aggregation of S-opsin (short-wavelength cone opsin, including uv opsin) is accompanied by the onset of cone degeneration through activating endoplasmic reticulum (ER) stress in a murine model^31^. Overall, impaired ocular phenotypes were caused by *blocs1s1*-overexpressing, but the impairment was milder than that in the knockdown groups above.

**Fig. 6.**
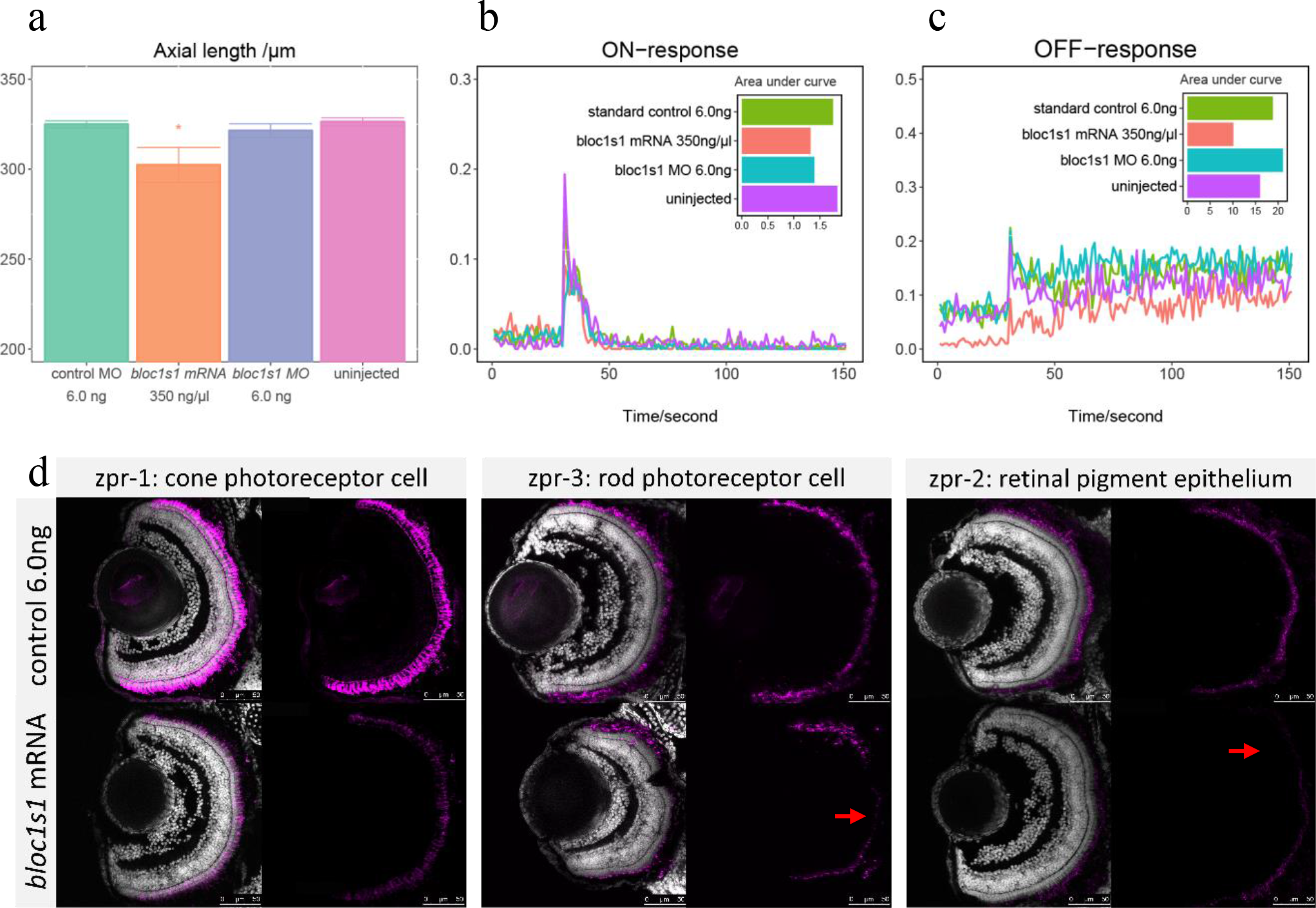
Phenotypes of *bloc1s1*-overexpression zebrafish larvae. (**a**) The bar plot of axial length. The *bloc1s1*-overexpression group showed a slight but significant reduction in ocular sizes. (**b, c**) The VMR testing for ON and OFF responses. The ON response was not apparently affected, but impairment of the OFF response was observed in the *bloc1s1* mRNA-injected group. (**d**) Immunostaining of *bloc1s1*-mRNA zebrafish retinae showed apparently low expression for zpr-1, zpr-2 and zpr-3.

Despite that the detrimental effects of overexpression of *bloc1s1* were moderate, the phenotypic results in zebrafish were in line with the estimated SMR effect in retina rather than that in blood. Here, the positive SMR effect means that increased expression level of a gene is associated with increased phenotype (or disease risk) and vice versa. The estimated SMR effect of *BLOC1S1* was in positive and those of *CNN2* and *SARM1* were negative, predicting that increased expression level of *BLOC1S1* and decreased expression levels of *CNN2* and *SARM1* are associated with increased AMD risk. In a word, the prediction from SMR and HEIDI analysis was validated in zebrafish that knockdown of *cnn2* and *sarm1* and overexpression of *bloc1s1* caused ocular abnormalities.

## Discussion

In this study, we aimed to shed light on the biological knowledge expected from integrated analysis of AMD GWAS and eQTL summary data. Initially, we identified 9 putative causal genes for AMD using the SMR and HEIDI methods based on a blood eQTL dataset (**Table 1**), and subsequently replicated 6 of the 9 genes using a retina eQTL dataset (**Table 1** and **Supplementary Figure 1**). We also tested the 9 genes in 48 other tissues using the GTEx eQTL data and showed that the across-tissue replication rate depended heavily on the size of replication sample (**Supplementary Figure 1**). We carried 4 of the 9 genes forward for functional assay in zebrafish and for the first time demonstrated the functional relevance of *CNN2*, *SARM1* and *BLOC1S1* to AMD. Injection of respective *cnn2* MO, *sarm1* MO and *bloc1s1* mRNA in zebrafish larvae exhibited different degrees of ocular defects (**Figures 2** and **6**), vision loss (**Figures 3** and **6**), relatively low transcription of essential photoreceptor and/or RPE genes (**Figure 4**) and RPE degeneration (**Figures 5** and **6**). Importantly, knockdown-induced phenotypes could be rescued by augmentation of corresponding mRNA (**Figure 3**), indicating the potential as targets to design new treatments. Finally, the phenotypic effects of alternating transcription of the prioritised genes in zebrafish were in line with the directions of estimated SMR effects in human retinas. All these results provide strong evidence supporting the potential role of *CNN2*, *SARM1* and *BLOC1S1* in AMD pathogenesis.

There is no denying that the study has some limitations. First, we used a large blood eQTL data set (*n* = 2,765) for discovery and a relatively small retina eQTL data set (*n* = 523) for replication, following the suggestion from a recent study^14^. This strategy would have missed genes with a cis-eQTL effect in retina but not in blood. However, we considered that if cis-eQTL effects are similar across tissues^18,24^, using a blood eQTL data set of large sample size could increase the power of gene discovery in comparison to using a retina eQTL data set of small sample size. In addition, blood is much more accessible than retina so that the growth of eQTL data from blood is expected to be faster than that from retina, suggesting that the power of our analysis strategy could be substantially improved in the future by leveraging blood eQTL data sets with sample sizes of orders of magnitude larger than that used in this study^35^.

Second, zebrafish is an ideal but not perfect animal model. AMD is an age-related disease. However, zebrafish larvae were studied instead of elder zebrafish because the most efficient duration of MO-mediated knockdown effects is the first two or three days of development, and efficiencies decrease later^36^, rendering it difficult to observe age-related morphological and functional changes. Generating a primate AMD model is the most scientifically valid since humans or primates are the only mammals with a macula and foveal centralis, where AMD manifests, but it is extremely challenging in particular with respect to costs and time scale of the experiment. Instead, an important role of the zebrafish models is to serve as an efficient screening test to narrow down the most plausible causal genes through our comprehensive evaluation approaches. And the robust ocular changes of the prioritised genes, even in larvae time, should not be underestimated. It is acknowledged that individual SNPs generally confer small effects on gene expression^11^, but MO interventions dramatically eliminate transcriptional levels, resulting in more severe and noticeable phenotypes. Moreover, accumulating evidence indicates that the origins of age-related disorders occur during foetal life^37,38^, and AMD should not be an exception.

Third, the genetic mechanisms of gene regulation and the relationship of gene regulation to AMD manifestation remain a mystery. Hypotheses of non-coding SNPs that influence gene expression include transcriptional, posttranscriptional, or posttranslational process^9^, such as non-coding RNA function or histone modification, allowing for more specific regulatory mechanistic studies. Of note, it is more important to identify the causal gene than the causal variant because the ultimate goal is to identify the causal gene, which can correspond to multiple causal variants. A mass of downstream research is required to understand the underlying molecular mechanisms. Intriguing clues include compelling biology such as inflammatory response and lipid metabolism and underlying overlapping pathophysiology with other age-related diseases for genes such as *CNN2*, *PILRB* and *ABCA7* that are also located at risk loci for late onset Alzheimer’s disease (AD) (**Supplementary Table 3** for the description of each prioritised gene).

In summary, we performed an integrative data analysis that efficiently pinpointed novel susceptibility genes for AMD, and demonstrated the functional relevance of *CNN2*, *SARM1* and *BLOC1S1* to the disease using zebrafish models. The gene discovery procedure, combining statistical analysis of large-scale data and experimental validation, can be applied to other complex disorders to fill the knowledge gaps between genetic variants and phenotypes.

## Online Methods

### Data used for the integrative analysis

The GWAS summary data used in this study were derived from the latest and largest AMD GWAS meta-analysis^5^ (see URLs section), consisting of 16,144 advanced AMD patients and 17,832 controls of predominantly European ancestry. The total number of SNPs was up to 12 million. The SNP effects were expressed as log odds ratios. Because the SNP allele frequency was not available, we estimated the allele frequencies using the UK Biobank data^16^.

The eQTL summary-level statistics were obtained from the CAGE data^11^, consisting of 36,778 transcription phenotypes and ~8 million SNPs on 2,765 peripheral blood samples (of predominantly European ancestry). Transcription levels were measured using Illumina gene expression arrays. For replication in the retina, the summary cis-eQTL data were obtained from 523 postmortem retinas with ~ 9 million SNPs and 15,124 gene expression traits^17^, available in the Eye Genotype Expression (EyeGex) database (URLs). For replication in other multiple tissues, we used the GTEx v7 data, containing a set of cis-eQTL summary data across 48 human tissues (URLs). Transcription levels in both EyeGex and GTEx were measured by RNA-seq. The eQTL effects in all the three data sets were expressed in standard deviation (SD) units of transcription levels.

### SMR and HEIDI test for pleiotropic association

SMR and HEIDI analyses were developed to identify genes whose expression levels were associated with a complex trait because of pleiotropy/causality (i.e., the trait and gene expression are associated due to the same set of causal variants at a locus)^12^. First, the SMR test takes the top associated cis-eQTL of the gene as an instrumental variable to test for association of a transcript (as an exposure) with AMD (as an outcome). An SMR estimate of the effect of gene expression on AMD is the ratio of the estimated effect of the instrument on a transcript (eQTL effect) and that on the disease (GWAS effect). The standard error (SE) of the SMR effect is computed using the Delta method, and the significance of the effect is assessed by the Wald test. To exclude the SMR associations due to linkage (i.e., the trait and gene expression are associated due to distinct set of causal variants in LD), the HEIDI analysis uses multiple SNPs in a cis-eQTL region to test against the null hypothesis that the trait is associated with gene expression because of the same set of underlying causal variants (pleiotropy/causality). Under the null, the SMR effects estimated using different eQTL SNPs in the cis region are expected to be the same. Significant heterogeneity in SMR effects detected at different SNPs in LD with the top associated cis-eQTL would be considered as linkage and rejected from the analysis.

### Morpholino, mRNA rescue and overexpression experiments

For the *cnn2* mRNA rescue experiment, we generated zebrafish cDNA from total RNA using RT-PCR on a full-length *cnn2* fragment. For the *sarm1* mRNA rescue and *bloc1s1* overexpression study, we attained specific cDNA from the pUC57 vector cloned into template cDNA sequences of *sarm1* or *bloc1s1*, which were obtained through oligoribonucleotide synthesis (Sangon Biotech, Shanghai, China). The amplification primers of *cnn2* included the forward (5’-TAATACGACTCACTATAGGGGCCACCATGTCTTCGCAG-3’) and the reverse (5’-TTAGTAATCTTGGCCGTCGTCCTGATAGC-3’); *sarm1* included the forward (5’-TAATACGACTCACTATAGGGGCCACCATGTTTTTGTCCCTCG-3’) and the reverse (5’-CTACTTCTTTTGTGGCTCTTTTTTGTCCG-3’); *bloc1s1*, included the forward (5’-TAATACGACTCACTATAGGGGCCACCATGCTCTCGCGG-3’) and the reverse (5’-TCATGTGGATGCCGGCTGGAC-3’); they all flank the T7 promoter sequence (5’-TAATACGACTCACTATAGGG-3’) and enhancing sequence (Kozak). PCR template DNA was purified using the QIAquick PCR Purification Kit (Qiagen, Germany). Capped and tailed full-length mRNA was then synthesized using an mMESSAGE mMACHINE™ T7 ULTRA Transcription Kit (Invitrogen, Carlsbad, CA) before purification using the RNeasy Mini Kit (Qiagen, Germany) following the manufacturer’s protocols. Rescue mRNAs were co-injected with MO into the one-cell stage embryos. For overexpression, corresponding mRNAs were injected into one- to two-cell stage embryos.

### Measurement of eye parameters and body length

The eye parameter and body length measurements of 3 days post-fertilisation (dpf) embryos were assessed using stereomicroscopy (SZX116, OLYMPUS, Japan). Pictures of the vertical and lateral view of each larva were recorded by a microscopic camera. Axial length, eye area, and body length were quantified by built-in software (OLYMPUS cellsens standard, version 1.14). For data collection, 10-15 larvae were included in each group; experiments were replicated three times. Student’s t-test was performed between controls and treatment conditions for each phenotype. Statistical differences were calculated and visualized by R software (version 3.5.3) and the ggplot2 package (version 3.1.0).

### Visual behaviour experiments

Visual motor response (VMR) of 5 dpf was measured using a Zebrabox (VMR machine ViewPoint 2.0, France) to evaluate lights-ON and lights-OFF responses. All larvae with different treatments were separately placed in a 96-well plate with adequate water to ensure free activities (12 larvae for each treatment). The Zebrabox protocol was set to apply: 1) dark adaption for 3 hrs; 2) ON light for 30 mins; 3) OFF light for 30 mins; and 4) repeat step 2 and step 3 three times. Motor activities were recorded every second. The duration of 150 s (30 s before and 120 s after light switching) was used to evaluate the visual motor activities of larvae. Aggregate data of three repetitions were compiled and visualized in the figures. The area under the curve, calculated by R package MESS (Version 1.0), was used to quantify the overall motor response during 150 s of light alterations. Twelve injected larvae for each group were randomly selected for experiments routinely conducted between 11:00 and 17:00.

### Real-time quantitative PCR

Zebrafish oculus was isolated from the control and experimental groups (n=30 pairs for each) at 3 dpf. Total RNA was then extracted with TRIzol reagent (Invitrogen Life Technologies, Carlsbad, CA, USA). RNA concentrations were determined using a NanoDrop instrument (NanoDrop Technologies, Thermo, US). Following the manufacturer’s instructions, purified RNA (500 ng) was used to generate cDNA using PrimeScript reverse transcriptase (TaKaRa, Dalian, China). Real-time quantitative PCR was performed through FastStart Universal SYBR Green Master (RocheApplied Science, Mannheim, Germany). Specific primers are provided in Supplementary Table 4. The relative expression levels were detected by the StepOne Plus TM Real-time PCR System (Life Technologies, Carlsbad, CA, USA) and calculated by the2-∆∆ CT method. The fold change was compared to the standard MO control. All experiments were performed in triplicate.

### Immunohistochemistry

For immunostaining purposes, morphants and control larvae at 3 dpf were fixed in 4% paraformaldehyde and were rinsed with 15% and 30% sucrose in PBS for dehydration. Zebrafish samples were then directly frozen in Richard-Allan Scientific™ Neg-50™. Cryosections of zebrafish eyes that were 18-mm were stained with zpr-1, zpr-2 and zpr-3 zebrafish-specific antibodies (1:400; provided by ZFIN) overnight at 4 °C. Alexa Fluor 594 anti-mouse secondary antibodies (1:400, provided by ZFIN) were used for incubation in blocking solution for 2 hrs at room temperature. DAPI (4,6-diamidino-2-phenylindole) was counterstained for the cell nucleus. Coverslips were mounted, and a confocal microscope (TCS SP8, Leica, Germany) was used to analyse gene expression and retinal architecture.

### URLs

AMD summary results: http://amdgenetics.org/

eQTL summary results: http://cnsgenomics.com/software/smr/#DataResource

SMR software: http://cnsgenomics.com/software/smr

EyeGex: https://gtexportal.org/home/datasets

Ensembl Orthologs: http://asia.ensembl.org/index.html

GeneCards Orthologs: https://www.genecards.org/

---

**Supplementary Figure 1:**
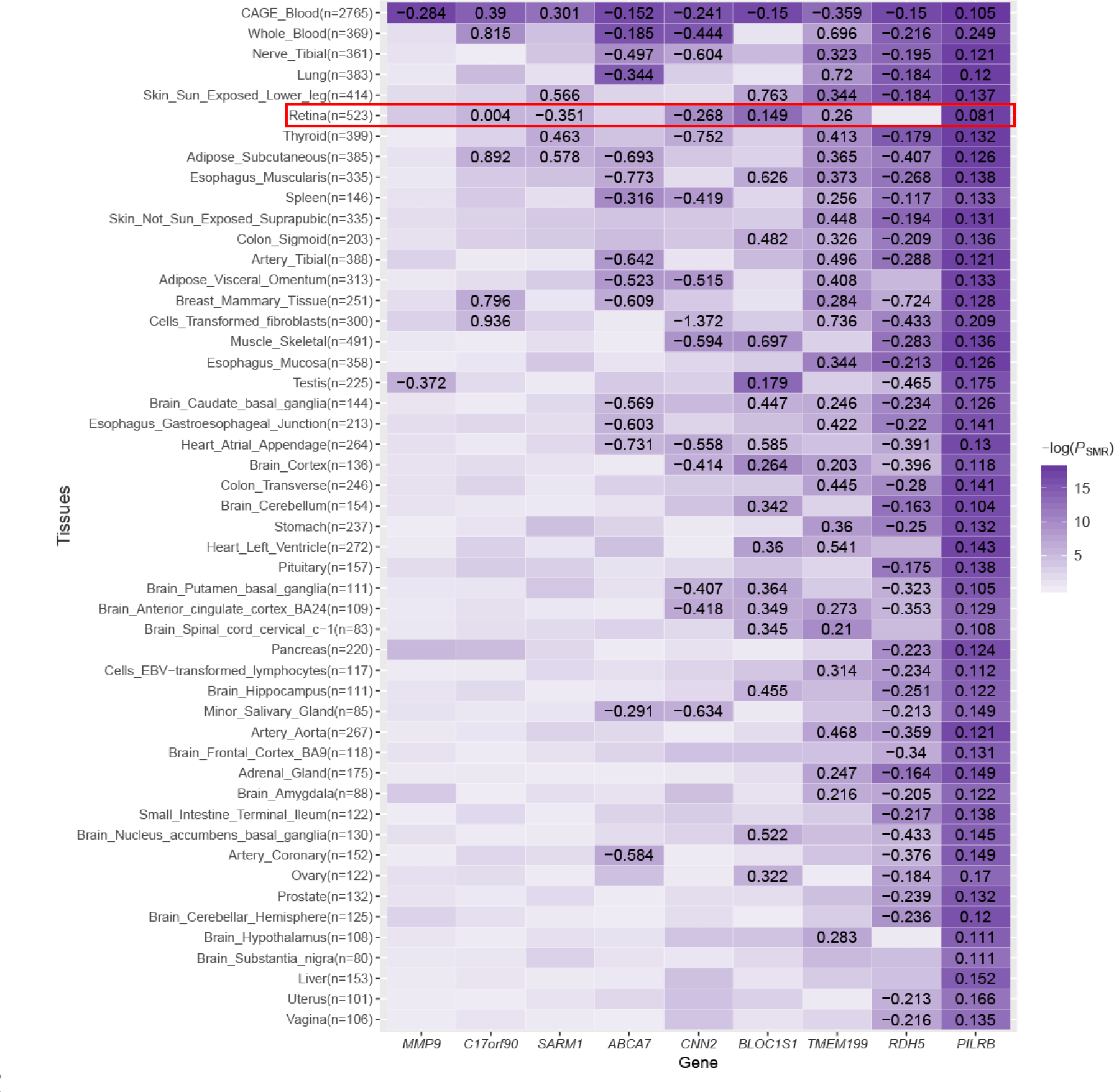
Heatmap of SMR results of the nine prioritised genes in multiple tissues. Each row represents a prioritised gene, and each column represents a tissue. −*log*(*P*-*value*_SMR_) is plotted in white-purple scale. The purple color indicates more significant and the white means less significant. Each tile with a number available indicates it reaches the significant threshold 5.6E−3 (correcting for 9 tests), with the number being the estimated SMR effect. Note that the overall mean SMR p-value is decreasing towards top and right. Replication in retina is highlighted by a red rectangle.

## Acknowledgements

This research was supported by the Natural Science Foundation of China (81522014; 81970838), National Key R&D Program of China (2017YFA0105300), and Zhejiang Provincial Natural Science Foundation of China (LQ17H120005), Ministry of Education 111 project (D16011), the Australian National Health and Medical Research Council (1113400) and the Australian Research Council (FT180100186). This study makes use of data from the UK Biobank (project ID: 21497). A full list of acknowledgments of this data set can be found in Supplementary Note 2.

