## Supplementary for "Towards the identification of causal genes for age-related macular degeneration"

**Supplementary Table 1: Putative functional genes for AMD from the SMR analysis in CAGE.** Detailed information of 16 genes (tagged by 21 probes) was gathered from SMR analysis at the genome-wide significance level.

| probeID | Chr | Gene | Probe bp | topSNP | topSNP bp | A1 | A2 | Freq | $\beta_{GWAS}$ | $P_{GWAS}$ | $\beta_{eQTL}$ | $P_{eQTL}$ | $\beta_{SMR}$ | $P_{SMR}$ | $P_{HEIDI}$ | $nsnp_{HEIDI}$ |
|---|---|---|---|---|---|---|---|---|---|---|---|---|---|---|---|---|
| ILMN_1660436 | 6 | *HSPA1A* | 31797709 | rs494620 | 31838713 | A | G | 0.384 | 0.079 | 3.71e-07 | 0.51 | 1.4e-74 | 0.155 | 9.72e-07 | 2.64e-08 | 20 |
| ILMN_1765532 | 6 | *RDBP* | 31919892 | rs550513 | 31920687 | T | C | 0.0979 | -0.534 | 9.46e-91 | -0.402 | 1.82e-16 | 1.33 | 2.45e-14 | 2.48e-07 | 20 |
| ILMN_1679520 | 6 | *AGPAT1* | 32136453 | rs41316748 | 32019512 | C | T | 0.0403 | 0.246 | 7.99e-10 | -0.72 | 1.29e-18 | -0.341 | 4.67e-07 | 4.8e-06 | 20 |
| ILMN_2044927 | 6 | *RNF5* | 32148208 | rs693906 | 31835164 | C | G | 0.157 | 0.153 | 7.05e-12 | 0.438 | 6.09e-27 | 0.349 | 7.45e-09 | 0.000454 | 20 |
| ILMN_1725170 | 6 | *CA425595* | 32624021 | rs9274614 | 32635846 | C | G | 0.241 | -0.141 | 6.5e-16 | 1.12 | 3.51e-267 | -0.126 | 3.5e-15 | 0.00326 | 20 |
| ILMN_1721636 | 7 | *TSC22D4* | 100064521 | rs7792525 | 99972122 | G | A | 0.188 | 0.112 | 1.09e-08 | 0.323 | 2.27e-20 | 0.347 | 1.16e-06 | 0.041 | 20 |
| ILMN_1688279 | 7 | *PVRIG* | 99818960 | rs6953580 | 99825275 | G | A | 0.216 | 0.090 | 1.1e-06 | -0.951 | 1.74e-180 | -0.0947 | 1.56e-06 | 0.051 | 20 |
| ILMN_1685534 | 7 | *PILRB* | 99947392 | rs7792525 | 99972122 | G | A | 0.188 | 0.112 | 1.09e-08 | 0.345 | 1.72e-23 | 0.325 | 7.01e-07 | 0.121 | 20 |
| ILMN_1723984 | 7 | *PILRB* | 99955692 | rs73401450 | 99981859 | C | G | 0.188 | 0.112 | 1.07e-08 | 1.07 | 3.28e-205 | 0.105 | 1.89e-08 | 0.183 | 20 |
| ILMN_1768754 | 7 | *PILRB* | 99965148 | rs61735533 | 99955866 | A | G | 0.188 | 0.111 | 1.6e-08 | 1.19 | 7.93e-255 | 0.093 | 2.48e-08 | 0.257 | 20 |
| ILMN_1807712 | 7 | *PILRB* | 99951516 | rs1964242 | 99976703 | A | G | 0.186 | 0.113 | 7.07e-09 | 0.649 | 4.41e-76 | 0.175 | 3.31e-08 | 0.317 | 20 |
| ILMN_1662839 | 10 | *PLEKHA1* | 124191568 | rs11200594 | 124139393 | C | T | 0.528 | -0.476 | 3.63e-211 | 0.385 | 1.78e-44 | -1.24 | 3e-37 | 1.07e-14 | 20 |
| ILMN_2394250 | 10 | *PLEKHA1* | 124189438 | rs10082476 | 124164654 | G | A | 0.244 | -0.316 | 2.31e-68 | 0.383 | 9.65e-30 | -0.826 | 2.01e-21 | 2.44e-06 | 20 |
| ILMN_1773395 | 12 | *BLOC1S1-RDH5* | 56118409 | rs56108400 | 56213297 | T | G | 0.242 | 0.103 | 2.36e-08 | -0.69 | 8.3e-82 | -0.15 | 8.31e-08 | 0.407 | 20 |

| | | | | | | | | | | | | | | | | |
|---|---|---|---|---|---|---|---|---|---|---|---|---|---|---|---|---|
| ILMN_2043615 | 17 | *C17orf90* | 79632146 | rs11150803 | 79621160 | A | C | 0.474 | 0.090 | 4.36e-09 | 0.539 | 6.77e-81 | 0.167 | 2.03e-08 | 0.0117 | 20 |
| ILMN_1746265 | 17 | *SARM1* | 26727880 | rs7212349 | 26733698 | T | C | 0.449 | -0.081 | 1.81e-07 | -0.268 | 1e-22 | 0.301 | 4.09e-06 | 0.128 | 20 |
| ILMN_1805131 | 17 | *C17orf90* | 79633554 | rs9910935 | 79613949 | T | C | 0.473 | 0.092 | 1.71e-09 | 0.237 | 1.13e-17 | 0.39 | 8.4e-07 | 0.139 | 20 |
| ILMN_1748481 | 17 | *TMEM199* | 26688817 | rs708100 | 26688663 | G | A | 0.488 | -0.086 | 2.5e-08 | 0.239 | 1.86e-18 | -0.359 | 2.56e-06 | 0.351 | 20 |
| ILMN_1743205 | 19 | *ABCA7* | 1065149 | rs3087680 | 1038289 | C | A | 0.111 | 0.171 | 4.57e-08 | -1.12 | 4.44e-136 | -0.152 | 9.33e-08 | 0.737 | 20 |
| ILMN_1708486 | 19 | *CNN2* | 1036186 | rs3087680 | 1038289 | C | A | 0.111 | 0.171 | 4.57e-08 | -0.709 | 3.26e-56 | -0.241 | 2.38e-07 | 0.76 | 20 |
| ILMN_1796316 | 20 | *MMP9* | 44644938 | rs3918261 | 44643592 | G | A | 0.143 | -0.134 | 1.02e-09 | 0.474 | 9.78e-34 | -0.284 | 4.98e-08 | 0.196 | 15 |

Chr represents chromosome; A1 is the effect allele; Freq is frequency of the effect allele in the reference sample.

**Supplementary Table 2: Orthologue similarity between human and zebrafish of the prioritised genes.** Data were extracted from the Ensembl and GeneCards websites.

| Gene | Species | Ensembl (%) | GeneCards(%) |
|---|---|---|---|
| *C17orf90* | Zebrafish | Lack of data | Lack of data |
| *pilrb* | Zebrafish | 20.26 | Lack of data |
| *abca7* | Zebrafish | 26.1 | 48 |
| *tmem199* | Zebrafish | 46.15 | 59.42 |
| *rdh5* | Zebrafish | 49.71 | 57.62 |
| *mmp9* | Zebrafish | 57.14 | 60.77 |
| *sarm1* | Zebrafish | 60.77 | 64.53 |
| *cnn2* | Zebrafish | 67.58 | 68.28 |
| *bloc1s1* | Zebrafish | 81.7 | 90 |

**Supplementary Table 3: Potential biological functions of the 9 putative AMD genes.** Shown are the results from manual literature search for the prioritised genes. Main biological functions involving in inflammatory response， angiogenesis, lipid metabolism and homeostasis, with partially overlapping associations to neurodegeneration diseases.

| Gene Name | Description |
|---|---|
| *ABCA7* | ATP-binding cassette sub-family A member 7. The mRNA has a dominant expression in myelo-lymphatic tissues[1] and microglia in brain[2]. It was suggested to play a role in macrophage transmembrane lipid transport[1]. Variants and epigenetic markers in ABCA7 have been reported significant association with Alzheimer's disease (AD)[3], also an age-related disease. Knockout of *Abca7* showed no obvious phenotypic abnormalities but with serum lipid alternation in young mice[4]. |
| *BLOC1S1* | Biogenesis of lysosome-related organelles complex 1 subunit 1. Mutation in the complexes results in Hermansky-Pudlak Syndrome, characterized by decreased pigmentation and lysosomal accumulation of ceroid lipofuscin, with also impaired vision[5]. |
| *C17orf90* | Also called *OXLD1*, Oxidoreductase Like Domain Containing 1. Relevant literature is unavailable. |
| *CNN2* | Calponin 2 is expressed in a broader range of tissues and a significant levels in macrophages. Deletion of *Cnn2* could accelerate macrophage migration and phagocytosis thus hinder the progress of atherosclerosis, a vascular inflammatory disease[6]. On the other hand, it is also demonstrated that *cnn2* MO zebrafish had cardiovascular defects[7]. Besides, alike *ABCA7*, *CNN2* resides in rs4147929 LD block, which is AD associated loci[8]. |
| *MMP9* | Matrix metallopeptidase 9, is specific to wet AMD among all AMD-associated variants[9]. In ophthalmology, *MMP9* participates in extracellular matrix remodeling and microvascular permeability during ocular angiogenesis in RPE and retinal microvascular endothelial cells[10]. It also acts as a bio-marker to identify inflammatory dry eye and ocular surface diseases[11]. Systemic pathological processes, like immunological diseases[12], cancer and its metastasis[13], cardiovascular[14]were found to associate *MMP9*. |
| *PILRB* | Paired immunoglobulin-like type 2 receptor beta, an activating immune receptor, distributes broadly across tissues and has a relative high expression in microglia[15]. Genetic variants in *PILRB* drive not only the association susceptibility of neurodegenerative diseases, such as AD, |

| | |
|---|---|
| | Parkinson's disease[16], but also altered expression of *PILRB*, where the expression was lower in AD cases compared with controls[8]. Additionally, rs61735533, labelling *PILRB* in the study, plays a larger role in European population (where it has a risk allele frequency of 18%-20%) than it does in Asian population (where is has a risk allele frequency of 3% in East Asian and 12% in South Asian), which is consistent with the prevalence difference of AMD in those population. |
| *RDH5* | Retinol Dehydrogenase 5. *RDH5* is one of causal genes of fundus albipunctatus, characterized with yellow and white lesion at RPE, with or without cone dystrophy[17]. Macular cone density was detected lower for fundus albipunctatus patients with *RDH5* mutations[18]. |
| *SARM1* | Sterile Alpha And TIR Motif Containing 1, a negative regulator of the Toll-like receptor signaling pathway in innate immunity and predominantly expressed in neurons[19]. Knockdown of Sarm1 negatively influences neuronal development and synaptic function[19,20]. It is also demonstrated loss-of-function of Sarm1 delays degeneration of injured axons by preventing ATP depletion[21,22]. |
| *TMEM199* | Transmembrane Protein 199. Deficiency of *TMEM199* leads to congenital disorders of glycosylation with hypercholesterolemia because of Golgi homeostasis disruption[23]. It is also involved in intracellular iron homeostasis required for endolysosomal acidification and lysosomal degradation[24]. |

**Supplementary Table 4: Primers for vital photoreceptors and RPE genes in real-time qPCR.**

| Gene | Forward Primer (5'-3') | Reverse Primer (5'-3') |
|---|---|---|
| *opn1lw2* *(red opsin)* | CCAACAGCAATAACACAAGGG | GCGACAACCACAAAGAACATC |
| *opn1mw1* *(green opsin)* | GGCTGTGTAATGGAGGGATTC | ATGGTTTGCGGAGAATTTGAAG |
| *opn1sw2* *(blue opsin)* | GGTTCCTTTCAGCACCATTG | AGAAGCCGAACACCATTACC |
| *opn1sw1* *(uv opsin)* | TCATTTTCTCCTACTCACAGCTC | CACAAAAGAGCCAACCATCAC |
| *rhodopsin* | AGTCCTGCCCAGACATCTAG | GTACTGTGGGTATTCGTATGGG |
| *rpe65a* | AGAGACGGGACGGTCTACAA | CCGTCATCCCAAAACTGTGC |

**Supplementary Figure 1: Heatmap of SMR results of the nine prioritised genes in multiple tissues.** Each row represents a prioritised gene, and each column represents a tissue. $-log(P\text{-}value_{\mathrm{SMR}})$ is plotted in white-purple scale. The purple color indicates more significant and the white means less significant. Each tile with a number available indicates it reaches the significant threshold 5.6E-3 (correcting for 9 tests), with the number being the estimated SMR effect. Note that the overall mean SMR p-value is decreasing towards top and right. Replication in retina is highlighted by a red rectangle.

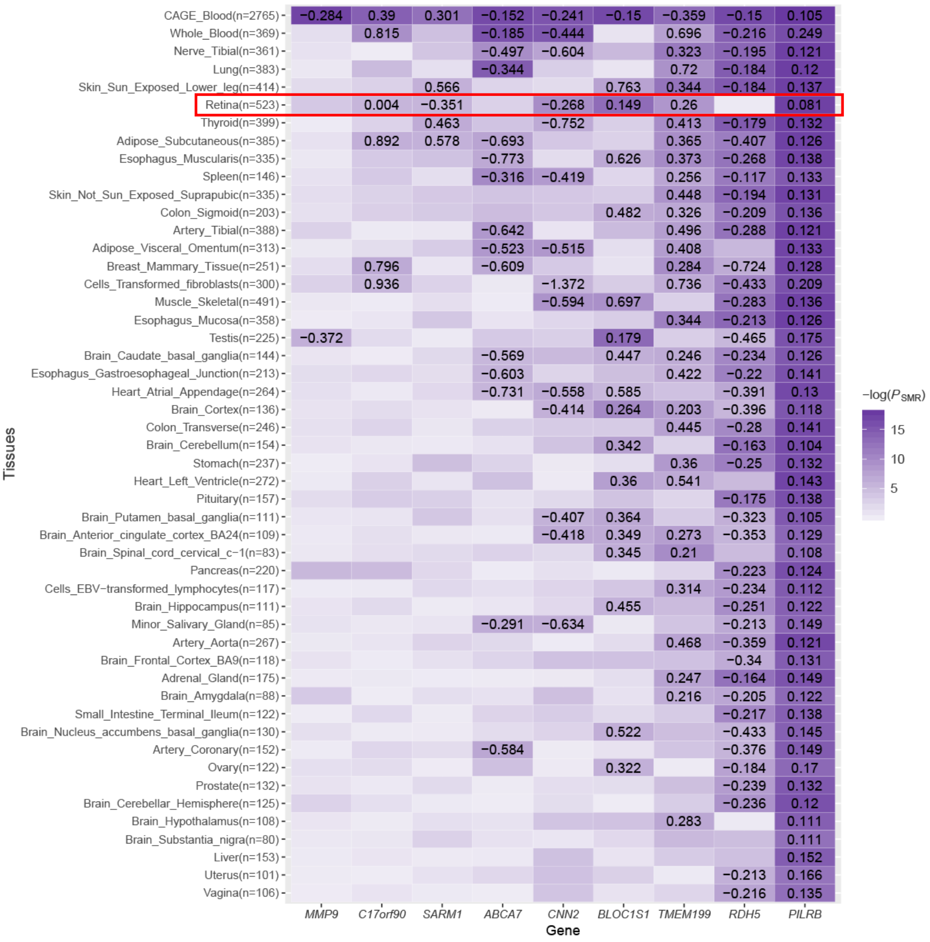

**Supplementary Figure 2: Prior experiments for morphology of *mmp9*-, *cnn2*-, *sarm1*- and *bloc1s1*-deficient zebrafish morphants.** All embryos were injected relevant MO at a dose of 6.0ng (N=10 for each group). (a, e) Lateral view of whole bodies. (b, f) Magnified lateral view of zebrafish eyeballs. (c, g) Vertical view of zebrafish eyeballs. (d, h) Quantification of body length, eye area, axial length and ratio of axial length and body length, respectively. Bar plot are shown in mean ± s.e.m. T-test was performed between each group with standard control. Significant reduction in axial length and eye area was observed in *cnn2*- and *sarm1*- deficient fishes.

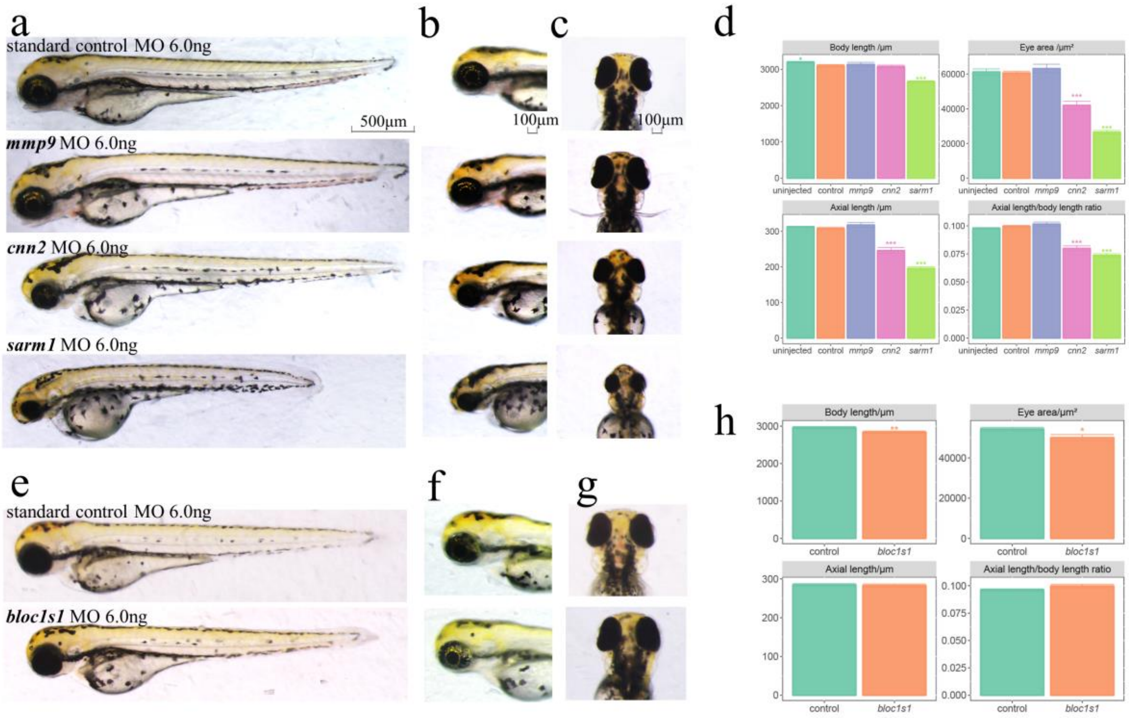

**Supplementary Figure 3: Dose-dependence of ocular phenotypes for *cnn2*-deficient zebrafish morphants.** The overall trends were declining as MO dose increased between 2.0-6.0 ng. N=10 for each group.

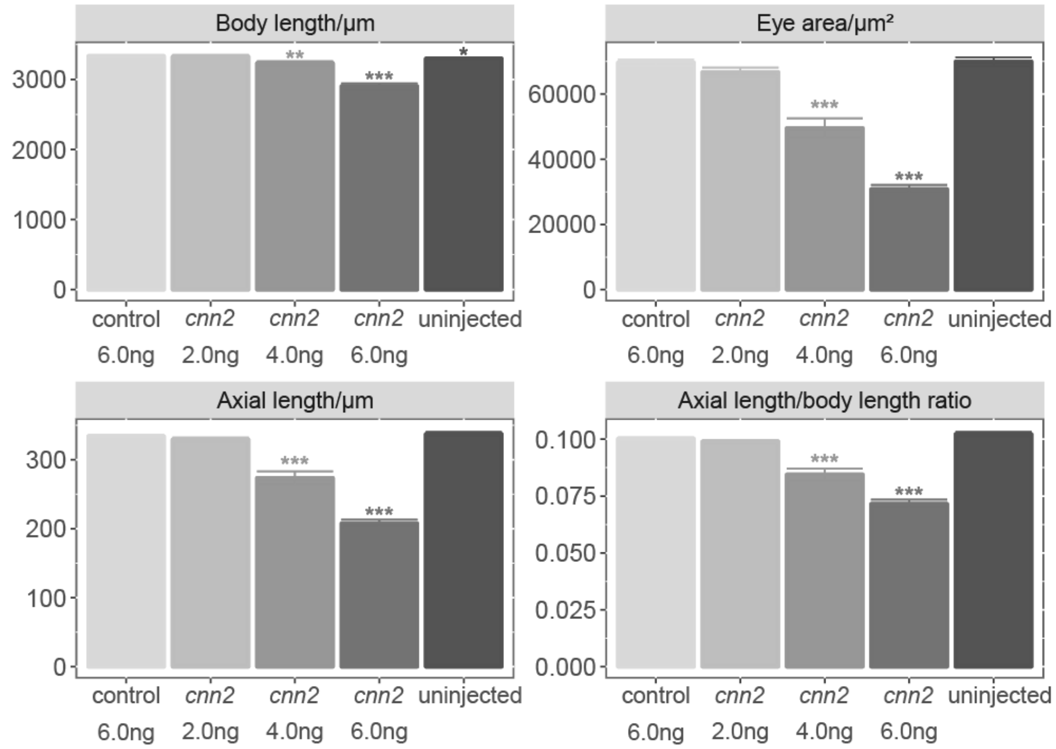

**Supplementary Figure 4: Dose-dependence of ocular phenotypes for *sarm1* deficient zebrafish morphants.** Decreased axial length and eye area occurred at a dose of 0.50 ng and decreased body length occurred at a dose of 0.75 ng. The overall trends were declining as MO dose increased before stayed stable at and more than 3.0 ng. N=10 for each group. The x-axis represents the dose of *sarm1* MO.

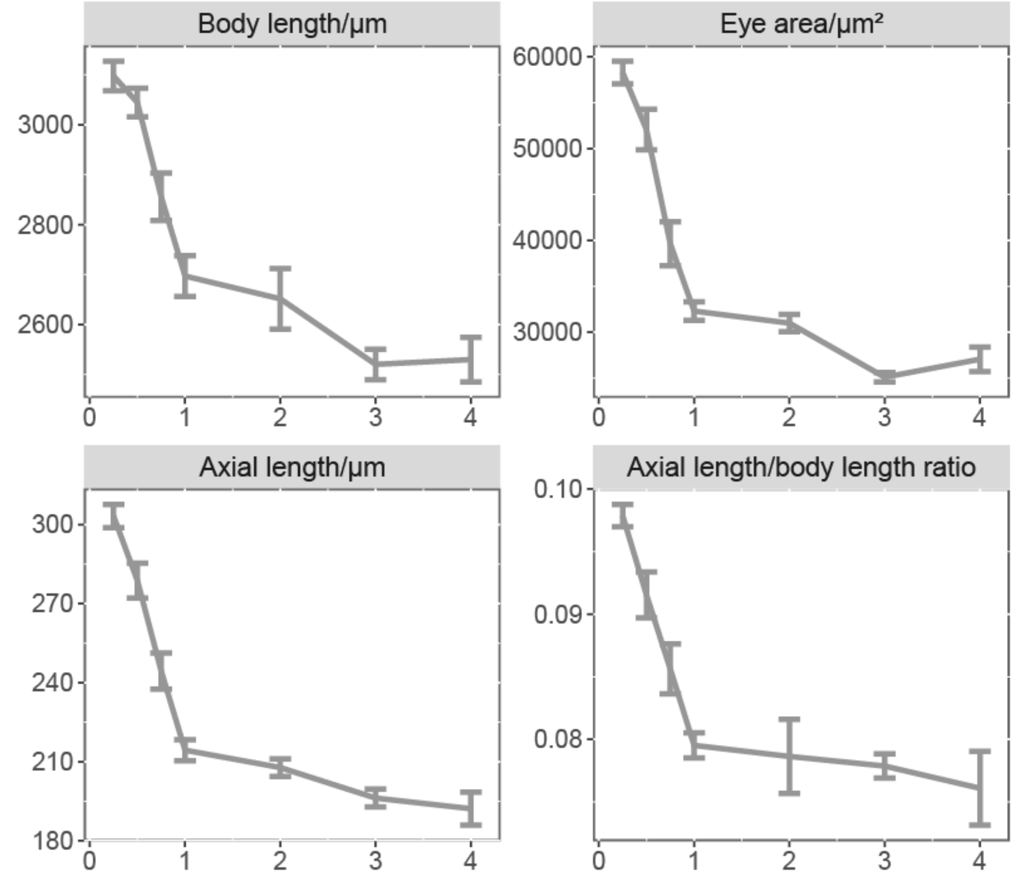

**Supplementary Figure 5: Pre-experiment for visual function of *cnn2* in zebrafish larvae.** Functional recovery relied on sufficient dose of injected mRNA. Partially rescue of visual function could be realized when injecting 200 ng/μl *cnn2* mRNA

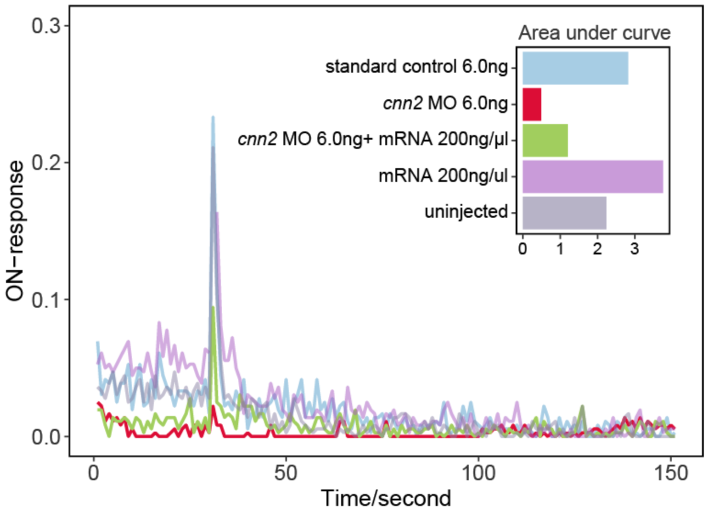


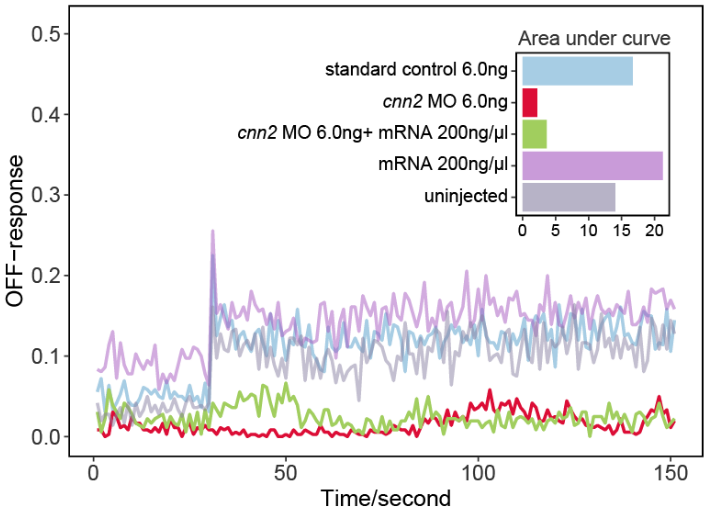

**Supplementary Figure 6: Retinal architecture of *sarm1*-deficient morphants.** There is a staining showing unimpaired rods in *sarm1* MO 1.0ng group.

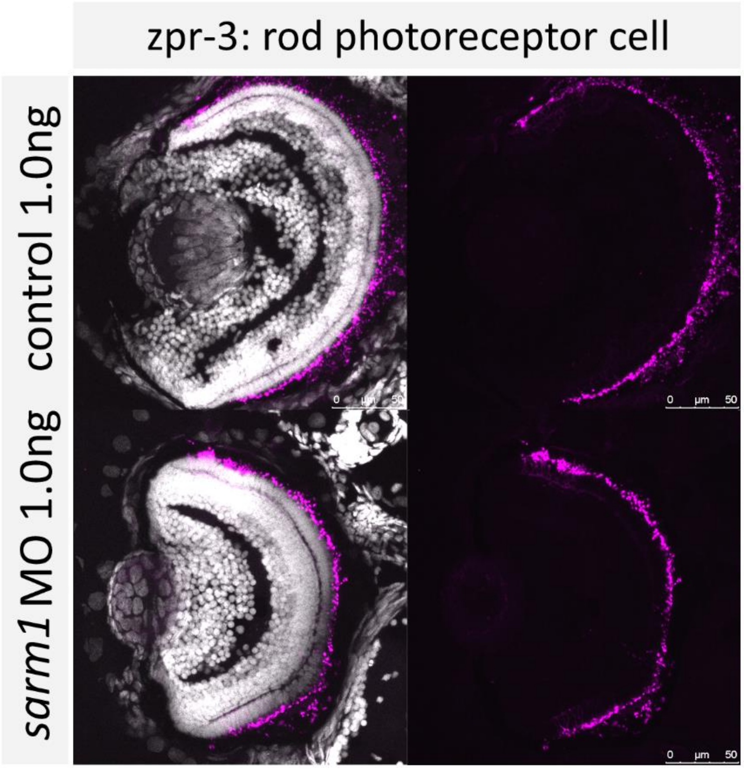

**Supplementary Figure 7: Prior experiment of *sarm1* 6.0ng MO group was observed severe lamination disruption.** Both cones and rods are also disrupted severely.

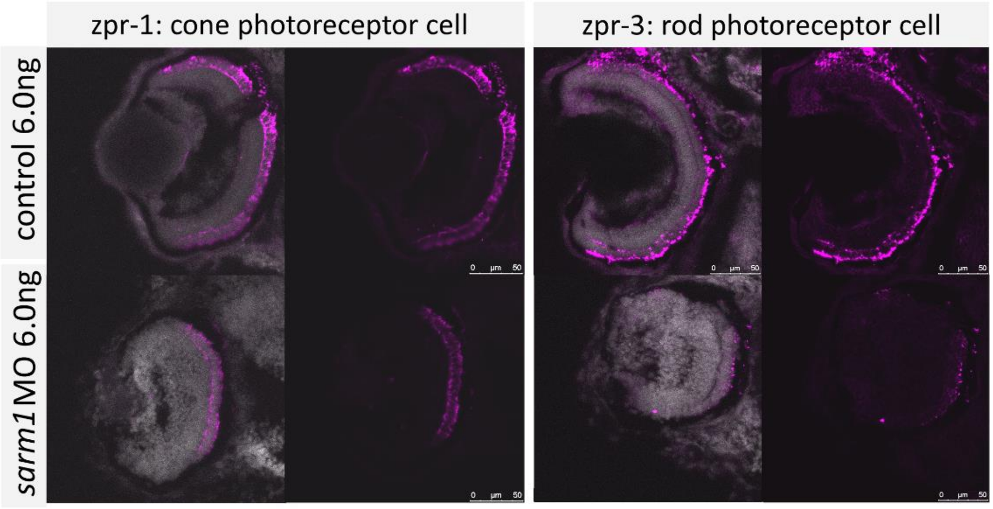

**Supplementary Figure 8: Staining repetition of *bloc1s1*-overexpressiong morphants.** Retinal architecture had no apparent difference in comparison to the standard control

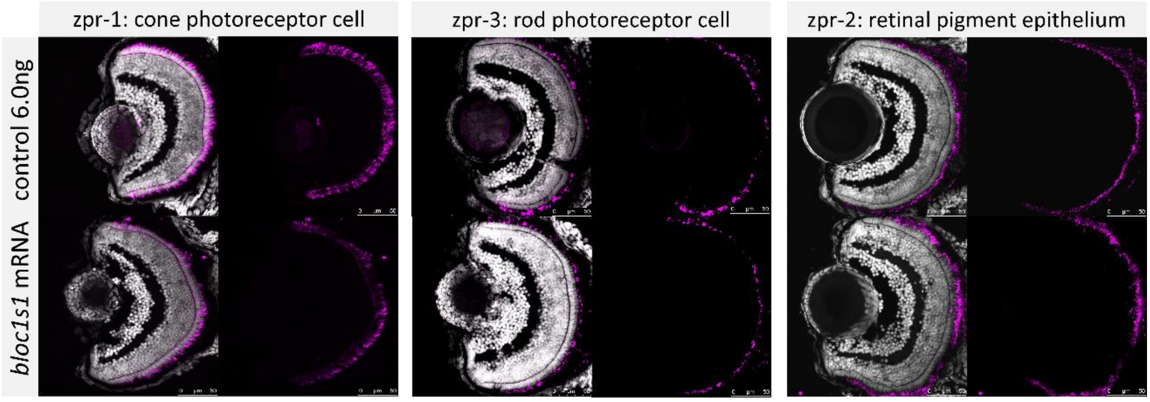

**Supplementary Figure 9: Real-time qPCR of photoreceptor and RPE genes in the *bloc1s1*-deficient and *bloc1s1*-overexpressiong zebrafish oculus.** The x-axis represents vital photoreceptors and RPE genes, and the y-axis represents the relative expression. Only rod gene for the *bloc1s1*-overexpressiong group was down-regulated (1.4-fold) and uv cone gene for both groups was up-regulated (1.3-2.3-fold) significantly but other retinal genes remained same. The bar plot is shown as the mean±s.e.m.

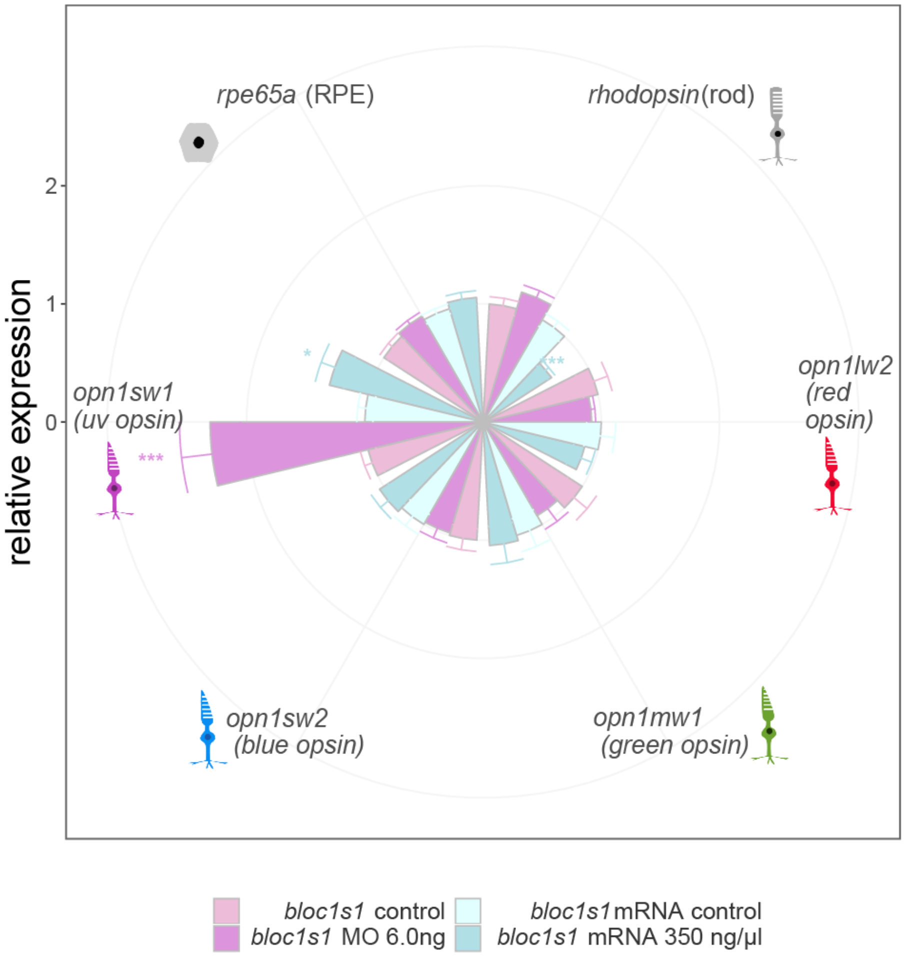

**Supplementary Note 1: Qualification of recovery degree of rescue experiments.**

$\frac{\overline{X}_{rescue} - \overline{X}_{MO}}{\overline{X}_{control} - \overline{X}_{MO}}$ , where $X$ is the axial length, eye area or motor activity in a certain condition.

**Supplementary Note 2: Acknowledgements**

**UKB:** This study has been conducted using UK Biobank resource under Application Number 21497. UK Biobank was established by the Wellcome Trust medical charity, Medical Research Council, Department of Health, Scottish Government and the Northwest Regional Development Agency. It has also had funding from the Welsh Assembly Government, British Heart Foundation and Diabetes UK.

### References – Supplementary Information:

1 Kaminski, W. E. *et al.* Identification of a Novel Human Sterol-Sensitive ATP-Binding Cassette Transporter (ABCA7). *Biochemical and Biophysical Research Communications* **273**, 532-538, doi:10.1006/bbrc.2000.2954 (2000).

2 Kim, W. S., Guillemin, G. J., Glaros, E. N., Lim, C. K. & Garner, B. Quantitation of ATP-binding cassette subfamily-A transporter gene expression in primary human brain cells. *Neuroreport* **17**, 891-896, doi:10.1097/01.wnr.0000221833.41340.cd (2006).

3 Rosenthal, S. L. & Kamboh, M. I. Late-Onset Alzheimer's Disease Genes and the Potentially Implicated Pathways. *Current genetic medicine reports* **2**, 85-101, doi:10.1007/s40142-014-0034-x (2014).

4 Kim, W. S. *et al.* Abca7 null mice retain normal macrophage phosphatidylcholine and cholesterol efflux activity despite alterations in adipose mass and serum cholesterol levels. *The Journal of biological chemistry* **280**, 3989-3995, doi:10.1074/jbc.M412602200 (2005).

5 Li, W. *et al.* Hermansky-Pudlak syndrome type 7 (HPS-7) results from mutant dysbindin, a member of the biogenesis of lysosome-related organelles complex 1 (BLOC-1). *Nature genetics* **35**, 84-89, doi:10.1038/ng1229 (2003).

6 Liu, R. & Jin, J. P. Deletion of calponin 2 in macrophages alters cytoskeleton-based functions and attenuates the development of atherosclerosis. *Journal of molecular and cellular cardiology* **99**, 87-99, doi:10.1016/j.yjmcc.2016.08.019 (2016).

7 Tang, J. *et al.* A critical role for calponin 2 in vascular development. *The Journal of biological chemistry* **281**, 6664-6672, doi:10.1074/jbc.M506991200 (2006).
8 Karch, C. M., Ezerskiy, L. A., Bertelsen, S. & Goate, A. M. Alzheimer's Disease Risk Polymorphisms Regulate Gene Expression in the ZCWPW1 and the CELF1 Loci. *PloS one* **11**, e0148717, doi:10.1371/journal.pone.0148717 (2016).
9 Fritsche, L. G. *et al.* A large genome-wide association study of age-related macular degeneration highlights contributions of rare and common variants. *Nature genetics* **48**, 134-143, doi:10.1038/ng.3448 (2016).
10 Das, A. *et al.* Human diabetic neovascular membranes contain high levels of urokinase and metalloproteinase enzymes. *Investigative ophthalmology & visual science* **40**, 809-813 (1999).
11 Kaufman, H. E. The practical detection of mmp-9 diagnoses ocular surface disease and may help prevent its complications. *Cornea* **32**, 211-216, doi:10.1097/ICO.0b013e3182541e9a (2013).
12 Gruber, B. L. *et al.* Markedly elevated serum MMP-9 (gelatinase B) levels in rheumatoid arthritis: a potentially useful laboratory marker. *Clinical immunology and immunopathology* **78**, 161-171 (1996).
13 Hiratsuka, S. *et al.* MMP9 induction by vascular endothelial growth factor receptor-1 is involved in lung-specific metastasis. *Cancer cell* **2**, 289-300 (2002).
14 Sakalihasan, N., Delvenne, P., Nusgens, B. V., Limet, R. & Lapiere, C. M. Activated forms of MMP2 and MMP9 in abdominal aortic aneurysms. *Journal of vascular surgery* **24**, 127-133 (1996).
15 Ryan, K. J. *et al.* A human microglia-like cellular model for assessing the effects of neurodegenerative disease gene variants. *Science translational medicine* **9**, doi:10.1126/scitranslmed.aai7635 (2017).
16 Li, Y. I., Wong, G., Humphrey, J. & Raj, T. Prioritizing Parkinson's disease genes using population-scale transcriptomic data. *Nature communications* **10**, 994, doi:10.1038/s41467-019-08912-9 (2019).
17 Nakamura, M., Hotta, Y., Tanikawa, A., Terasaki, H. & Miyake, Y. A high association with cone dystrophy in Fundus albipunctatus caused by mutations

of the RDH5 gene. *Investigative ophthalmology & visual science* **41**, 3925-3932 (2000).

18 Makiyama, Y. *et al.* Cone abnormalities in fundus albipunctatus associated with RDH5 mutations assessed using adaptive optics scanning laser ophthalmoscopy. *American journal of ophthalmology* **157**, 558-570.e551-554, doi:10.1016/j.ajo.2013.10.021 (2014).

19 Lin, C. W., Chen, C. Y., Cheng, S. J., Hu, H. T. & Hsueh, Y. P. Sarm1 deficiency impairs synaptic function and leads to behavioral deficits, which can be ameliorated by an mGluR allosteric modulator. *Frontiers in cellular neuroscience* **8**, 87, doi:10.3389/fncel.2014.00087 (2014).

20 Lin, C. W. & Hsueh, Y. P. Sarm1, a neuronal inflammatory regulator, controls social interaction, associative memory and cognitive flexibility in mice. *Brain, behavior, and immunity* **37**, 142-151, doi:10.1016/j.bbi.2013.12.002 (2014).

21 Yang, J. *et al.* Pathological axonal death through a MAPK cascade that triggers a local energy deficit. *Cell* **160**, 161-176, doi:10.1016/j.cell.2014.11.053 (2015).

22 Osterloh, J. M. *et al.* dSarm/Sarm1 is required for activation of an injury-induced axon death pathway. *Science (New York, N.Y.)* **337**, 481-484, doi:10.1126/science.1223899 (2012).

23 Jansen, J. C. *et al.* TMEM199 Deficiency Is a Disorder of Golgi Homeostasis Characterized by Elevated Aminotransferases, Alkaline Phosphatase, and Cholesterol and Abnormal Glycosylation. *American journal of human genetics* **98**, 322-330, doi:10.1016/j.ajhg.2015.12.011 (2016).

24 Miles, A. L., Burr, S. P., Grice, G. L. & Nathan, J. A. The vacuolar-ATPase complex and assembly factors, TMEM199 and CCDC115, control HIF1alpha prolyl hydroxylation by regulating cellular iron levels. *eLife* **6**, doi:10.7554/eLife.22693 (2017).
